# A trans-synaptic IgLON adhesion molecular complex directly contacts and clusters a nicotinic receptor

**DOI:** 10.1101/2024.09.05.611427

**Authors:** Morgane Mialon, Liubov Patrash, Alexis Weinreb, Engin Özkan, Jean-Louis Bessereau, Berangere Pinan-Lucarre

**Author notes:** Co-last, co-corresponding authors (Jean-Louis Bessereau, Berangere Pinan-Lucarre).

## Abstract

The localization and clustering of neurotransmitter receptors at appropriate postsynaptic sites is a key step in the control of synaptic transmission. Here, we identify a novel paradigm for the synaptic localization of an ionotropic acetylcholine receptor (AChR) based on the direct interaction of its extracellular domain with a cell adhesion molecule of the IgLON family. Our results show that RIG-5 and ZIG-8, which encode the sole IgLONs in *C. elegans,* are tethered in the pre- and postsynaptic membranes, respectively, and interact *in vivo* through their first immunoglobulin-like (Ig) domains. In addition, ZIG-8 traps ACR-16 via a direct *cis-*interaction between the ZIG-8 Ig2 domain and the base of the large extracellular AChR domain. Such mechanism has never been reported, but all these molecules are conserved during evolution. Similar interactions may directly couple Ig superfamily adhesion molecules and members of the large family of Cys-loop ionotropic receptors, including AChRs, in the mammalian nervous system, and may be relevant in the context of IgLON-associated brain diseases.

## INTRODUCTION

Chemical neurotransmission relies on the precise anchoring and clustering of postsynaptic receptors directly opposite specific neurotransmitter release sites. This process is tightly controlled by cell surface and secreted molecules that interconnect the molecular components of the synapse and ensure accurate alignment between the presynaptic and postsynaptic domains^1^. However, the diversity of molecular players that govern receptor clustering is becoming increasingly complex, raising new questions about the mechanisms regulating synapse formation and homeostasis.

One prevailing mechanism of postsynaptic receptor clustering involves the recruitment of intracellular protein scaffolds, composed of multimodular proteins of the postsynaptic density that multimerize and anchor postsynaptic receptors through intracellular binding. Synaptic transmembrane proteins localize these scaffolds, resulting in the formation of large molecular assemblies that span the postsynaptic membranes. In the mammalian brain, prototypical examples of these cytoplasmic scaffolds include PSD95 (PostSynaptic Density protein 95), PSD97, Homer and Shank at excitatory synapses, and Gephyrin and Collibistin at inhibitory synapses^2,3^.

More recently, an increasing number of components of the synaptic extracellular matrix have been demonstrated to serve as scaffolding molecules and help recruit synaptic receptors through extracellular interactions (reviewed in^4^). For example, secreted neuronal pentraxins (Nptx), and the transmembrane-bound neuronal pentraxin receptor (Nptxr) associate with AMPARs via their pentraxin domains *in vitro,* and cluster AMPARs *in vivo*, likely through self-multimerization^5,6^. Similarly, Cerebellin 1 (Cbln1), Cerebellin 4 (Cbln4) and the C1ql2-3 complement-related molecules are secreted from presynaptic terminals and bind the atypical GluD1-2 glutamate receptor delta subunits and the GluK2-4 kainate receptor subunits, respectively^7–9^. Both Cerebellins and C1ql2-3 are part of large complexes that are stabilized through their own oligomerization and bridge pre- and postsynaptic components.

Interestingly, both mechanisms can be used at the same synapse. In the nematode *C. elegans*, the extracellular matrix MADD-4/Punctin is secreted by motoneurons and localizes two different ionotropic acetylcholine receptors (AChRs) at excitatory neuromuscular junctions (NMJs) via two distinct mechanisms. The levamisole-sensitive acetylcholine receptors (L-AChRs) are heteropentameric AChRs that are activated by the nematode-specific cholinergic agonist levamisole^10^. L-AChRs interact with a *bona fide* extracellular complex consisting of two secreted proteins, LEV-9 and OIG-4, and the ectodomain of a transmembrane protein, LEV-10; all expressed by muscle cells^11–13^. These proteins form L-AChR-containing microclusters that are recruited and stabilized at synapses, most likely through direct interaction with Punctin in the synaptic cleft^14^. The nicotine-sensitive AChRs (N-AChRs) are homopentameric receptors composed of the ACR-16 subunit, the ortholog of the mammalian α7 AChR subunit^15^. Their synaptic localization depends on their interaction with an intracellular scaffold. At the cholinergic NMJ, Punctin activates the netrin receptor UNC-40/DCC and localizes the transmembrane heparan sulfate proteoglycan receptor Syndecan^16–18^. This dual signaling triggers the subsynaptic recruitment of the scaffolding proteins LIN-2/CASK and FRM-3/FARP1-2, which physically bind the large intracellular loop of the ACR-16 receptor and localize it at the NMJ.

In *C. elegans*, acetylcholine is not only used at NMJs but is the major neurotransmitter throughout the nervous system. Neurons express a large diversity of AChRs. In particular, the gene *acr-16* is readily expressed in several neurons but the mechanisms controlling its subcellular localization are unknown. To identify novel mechanisms of synapse organization, we performed an innovative genetic analysis of ACR-16-containing neuron-neuron synapses. Here we describe a critical role of two cell adhesion molecules, namely RIG-5 (neuRonal IG CAM-5) and ZIG-8 (Zwei IG domain protein-8), which are the sole IgLON orthologs in *C. elegans*. IgLONs are Ig-domain rich extracellular proteins that have often been suggested to act as synaptic adhesion molecules across phyla, but their actual synaptic functions have remained largely unexplored. We found that RIG-5 and ZIG-8 are specifically localized to pre- and postsynaptic membranes, respectively, and interact trans-synaptically through their N-terminal Ig1 domains. Most notably, we demonstrated that ZIG-8 recruits AChRs through a direct physical *cis*-interaction involving its Ig2 domain and the base of the AChR ectodomain. This is an unprecedented mechanism controlling the synaptic localization of an AChR through a direct extracellular interaction with synaptic adhesion molecules.

## RESULTS

### The ACR-16 acetylcholine receptor forms neuronal clusters in the ventral nerve cord

To visualize ACR-16 in living animals, we used a knock-in allele expressing a Scarlet-tagged version of ACR-16 **(Supplementary Figure 1A)**^16^. As previously described, ACR-16-Scarlet appeared at NMJs as characteristic spreading clusters along the ventral and dorsal sides of the animals. In addition, we also visualized large, round and sharply defined clusters in the ventral nerve cord **(Supplementary Figure 1B)**. These two types of clusters followed distinct tracks. Interestingly, transcriptomic data from single-neuron analyses indicate that *acr-16* is expressed in several neurons within the ventral nerve cord **(Supplementary Figure 1C)**^19^. This nerve structure is a major neuropil consisting of a bundle of approximately 40 neurites and hundreds of en passant synapses that extends from the head to the tail **(Figure 1A)**^20^. To investigate whether the round ACR-16 clusters are formed in neurons, we performed tissue-specific degradation of ACR-16 by utilizing the Auxin-Inducible Degradation (AID) system^21^. We introduced the AID cassette in the tagged *acr-16::Scarlet* locus using CRISPR/Cas9 and triggered proteasomal degradation of ACR-16-AID-Scarlet in body wall muscle cells and neurons **(Supplementary Figure 1A)**. Degrading ACR-16-AID-Scarlet in body wall muscle cells eliminated the spreading clusters, confirming their presence at NMJs **(Figure 1B)**. Conversely, degrading ACR-16-AID-Scarlet in neurons caused the disappearance of the round clusters, indicating their neuronal origin. Simultaneous degradation in both neurons and muscle cells completely removed ACR-16-AID-Scarlet, demonstrating highly efficient degradation. For this study, we focused primarily on the neuronal clusters of ACR-16-AID-Scarlet and, unless otherwise specified, we induced ACR-16-AID-Scarlet degradation specifically in body wall muscle cells.

**Figure 1.**
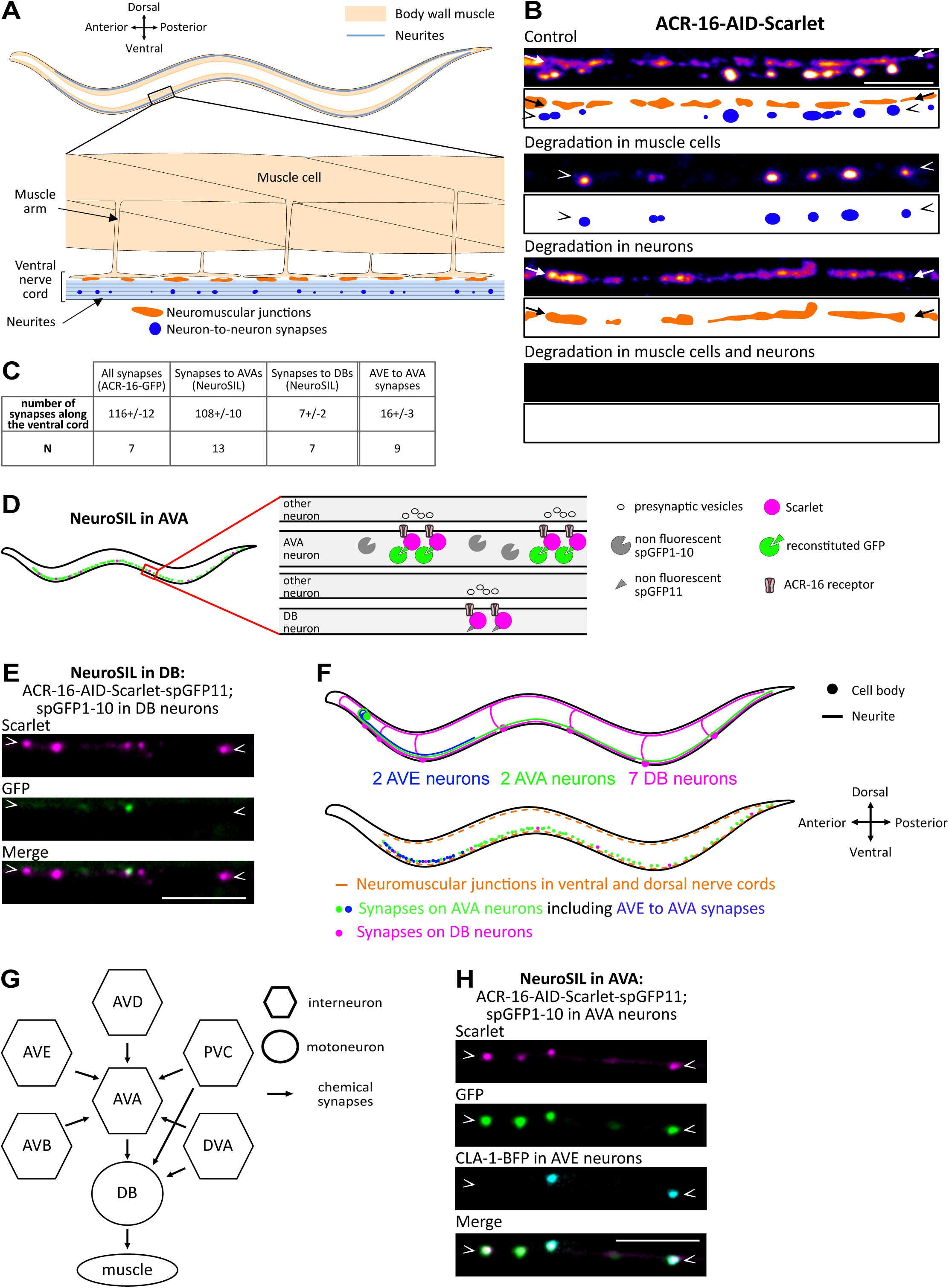
ACR-16 acetylcholine receptors form synaptic clusters in neurons of the ventral nerve cord. **(A)** Schematic of an adult *C. elegans* with a zoom-in showing body wall muscle cells and the ventral nerve cord, including NMJs and neuron-neuron synapses. **(B)** Spinning disk microscopy images and schematics of ACR-16-AID-Scarlet along the ventral cord at NMJs (orange dots, arrows) and neuron-neuron synapses (blue dots, arrowheads) under control conditions and following auxin-induced degradation in muscle cells, neurons and both. In each figure panel, arrowheads mark the location of neurites expressing ACR-16. **(C)** Number of ACR-16 clusters along the entire ventral nerve cord (ACR-16-GFP), in AVA and in DB neurons, and at AVE to AVA synapses (colocalizing with a CLA-1-BFP presynaptic marker expressed in AVE neurons). **(D)** Schematic of the NeuroSIL strategy for AVA neurons in the ventral nerve cord using the ACR-16-AID-Scarlet-spGFP11 strain. Upon auxin-induced degradation in muscle, all neuronal ACR-16 clusters are marked with Scarlet fluorescence (magenta), while ACR-16 clusters formed in AVAs are also detectable following neuron-specific co-expression of a spGFP1-10 moiety (green). **(E)** NeuroSIL images in DB neurons. **(F)** Schematic of an adult *C. elegans* from a lateral view showing the indicated neurons, and synapses with ACR-16. **(G)** Connectivity diagram of AVA neurons, DB neurons and their presynaptic partners at chemical synapses in the ventral nerve cord, according to White *et al.*, 1986^20^. **(H)** NeuroSIL images in AVA neurons showing Scarlet fluorescence and GFP reconstruction, along with presynaptic expression of CLA-1-BFP in AVE neurons. Scale bars: 5 µm (B, E and H).

### The ACR-16 acetylcholine receptor forms postsynaptic clusters at AVE to AVA synapses

To identify the neurons of the ventral nerve cord in which ACR-16 clusters, we first examined single-neuron transcriptomic data and identified four classes of *acr-16* expressing neurons that potentially synapse in the ventral nerve cord based on neuronal wiring data **(Supplementary Figure 1C)**^19,20^. In parallel, we constructed a fluorescent transcriptional reporter driven by a 5.8 kb upstream promoter sequence of *acr-16* **(Supplementary Figure 1A)**. Its expression pattern was analyzed in the NeuroPAL strain, which facilitates neuron identification through unique color labels and positions^22^. We confirmed *acr-16* expression in two classes of neurons that synapse in the ventral nerve cord: the AVA, a pair of interneurons that control locomotion, and the DB-type of cholinergic motoneurons **(Supplementary Figure 1C, D)**.

Using fluorescently-tagged ACR-16, we counted on average 116 neuronal clusters along the ventral cord **(Figure 1C, Supplementary Figure 1E)**. To specifically evaluate the number of clusters formed in the AVA and DB neurons, we developed a versatile strategy based on split GFP reconstitution in the cytoplasm of specific neurons, which we named NeuroSIL (Neuron-type Specific Illumination, **Figure 1D**)^23^. We inserted three copies of the spGFP11 sequence into the *acr-16::aid::Scarlet* locus via CRISPR/Cas9 (*acr-16::aid::Scarlet::spgfp11*) and expressed the complementary spGFP1-10 moiety in the AVA or in the DB neurons **(Supplementary Figure 1A)**^24^. Using NeuroSIL, the red Scarlet fluorescence labels the full complement of ACR-16 clusters, while the GFP fluorescence is reconstituted only in the neurons expressing spGFP1-10. In the AVA neurons, NeuroSIL revealed on average 108 ACR-16-Scarlet and GFP positive clusters **(Figure 1C, Supplementary Figure 1F)**. Because some clusters showed only Scarlet fluorescence and thus were not formed in the AVAs, we also tested NeuroSIL in the DBs **(Figure 1C, E)**. This experiment revealed on average 7 GFP+Scarlet clusters in the DB neurons. We concluded that the DBs and AVAs are the two neuron types in which ACR-16 form clusters, with the latter contributing to the majority of these clusters **(Figure 1F)**.

The AVA and DB neurons receive numerous synaptic inputs from several cholinergic neurons in the ventral nerve cord, as evidenced by electron microscopy **(Figure 1G)**^20,25^. Notably, the AVE interneurons establish synapses with the AVA neurons in the first quarter of the ventral nerve cord **(Figure 1F)**. To investigate the pattern of presynaptic sites in AVE neurons, we built an active zone reporter using CLA-1/Clarinet^26^. Our results revealed the juxtaposition of approximately 16 active zones and ACR-16-AID-Scarlet clusters in the first quarter of the ventral nerve cord, indicating that ACR-16 forms postsynaptic clusters at AVE to AVA synapses **(Figure 1C, G, H)**. Using the same methodology, we examined a presynaptic AVB reporter, as AVB neurons have been reported to make approximately 27 synapses on AVA by electron microscopy^20,25^. However, we did not detect strong juxtaposition of the pre- and postsynaptic signals in this context, suggesting that ACR-16 is not present at AVB to AVA synapses **(Supplementary Figure 1G)**. Owing to the lack of specific promoters for certain neuron types, such as AVD, which forms many synapses on AVA, we could not probe every potential presynaptic neuron of the AVAs and DBs^20,25^. Nevertheless, our data identify that ACR-16 form postsynaptic clusters at least at AVE to AVA synapses.

### The IgLON adhesion molecules RIG-5 and ZIG-8 specifically associate with ACR-16 clusters

To elucidate the molecular mechanisms controlling ACR-16 synaptic localization, we performed a genetic screen for mutants with altered patterns of ACR-16-AID-Scarlet clustering, and identified several mutations in the *rig-5* and *zig-8* genes **(Supplementary Figure 2A, B, Supplementary Table S2)**. These genes encode orthologs of mammalian IgLON cell adhesion molecules, also known as DIPs (Dpr Interacting Protein) and Dprs (Defective Proboscis extension Response) in *Drosophila*^27,28^. RIG-5 and ZIG-8 are composed of three and two extracellular Ig-like domains, respectively, such as DIPs and Dprs, and are tethered to the plasma membrane via glycosylphosphatidylinositol (GPI) anchoring^29,30^. Using CRISPR/Cas9, we engineered BFP-RIG-5 and GFP-ZIG-8 translational reporters by tagging them with fluorescent proteins immediately after their signal peptides. Strikingly, both BFP-RIG-5 and GFP-ZIG-8 exhibited round puncta that overlapped extensively with ACR-16-AID-Scarlet clusters along the ventral nerve cord **(Figure 2A)**. Despite the abundance of cholinergic synapses in the ventral nerve cord, BFP-RIG-5 and GFP-ZIG-8 were only present along with ACR-16-AID-Scarlet, and not at other types of synapses **(Figure 2A, Supplementary Figure 1H)**. This evidence strongly supports the conclusion that RIG-5 and ZIG-8 are cell adhesion molecules specifically associated with ACR-16 clusters.

**Figure 2.**
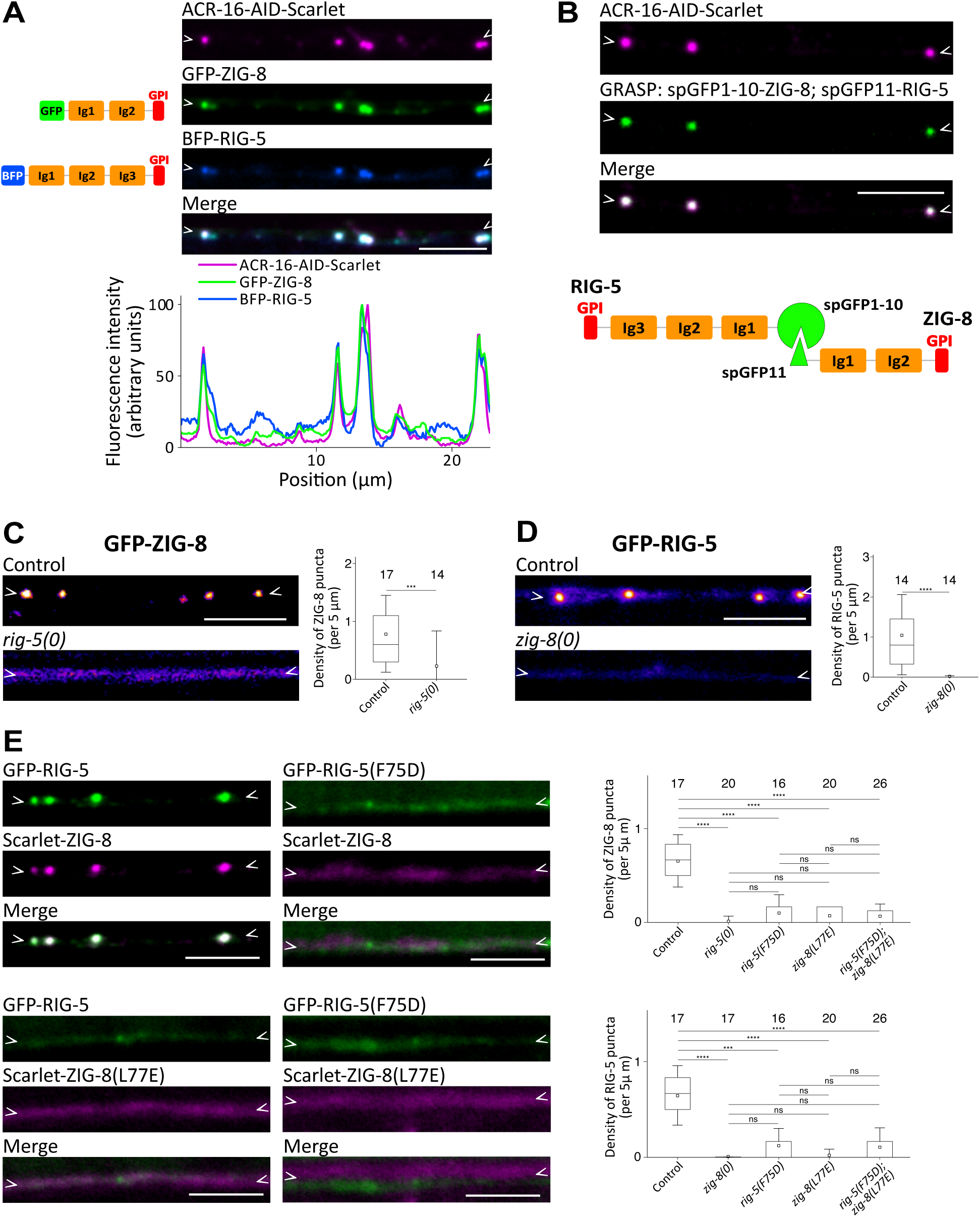
RIG-5 and ZIG-8 interact at synapses that express ACR-16. **(A)** Knock-in reporters of BFP-RIG-5 and GFP-ZIG-8 show strong colocalization with neuronal ACR-16-AID-Scarlet upon auxin-induced degradation in muscle. The fluorescence intensity profile is shown below. Left: schematic of BFP-RIG-5 and GFP-ZIG-8 functional domains. **(B)** Neuronal ACR-16-AID-Scarlet clusters observed following auxin-mediated degradation of ACR-16 in muscle, and GRASP between RIG-5 and ZIG-8 as indicated. **(C-D)** Fluorescence images and puncta density of GFP-ZIG-8 in a control strain and in the *rig-5(0)* mutant (C), and of BFP-RIG-5 in a control strain and in the *zig-8(0)* mutant (D). **(E)** GFP-RIG-5 and Scarlet-ZIG-8 were observed in control condition, and using strains bearing missense mutations that prevent the RIG-5—ZIG-8 interaction: *gfp::rig-5(F75D)* and *Scarlet*::*zig-8(L77E)* single mutants, and the *gfp::rig-5(F75D)*; *Scarlet::zig-8(L77E)* double mutant. The density of RIG-5 and ZIG-8 puncta is shown. Data are presented as boxplots showing lower and upper quartiles (box), mean (square), median (center line) and standard deviation (whiskers); the number of worms is indicated for each condition; Wilcoxon Mann-Whitney test (C, D), Kruskal-Wallis and Dunn’s test (E); ns: non-significant, ***<0.0005, ****<0.00005. Scale bars: 5 µm.

### RIG-5 and ZIG-8 control each other’s localization through interactions between their N-terminal Ig1 domains

A peculiar feature of IgLONs is their involvement in multiple homo- and heterophilic interactions between members of the family, in *trans* across different cells^28,31^. For several dimers, the structural basis of these interactions has been solved through crystallography, revealing a central hydrophobic patch of amino acids in the first N-terminal Ig1 domains as crucial for binding^28,32,33^. RIG-5 and ZIG-8 were demonstrated to form heterodimers by interacting in their Ig1 domains, as evidenced by crystallography and binding analysis, akin to mammalian and *Drosophila* IgLONs^29^. Given the *in vitro* interaction between RIG-5 and ZIG-8, we tested whether this interaction applies *in vivo* in the synaptic cleft. Firstly, we performed a GFP Reconstruction Across Synaptic Partners (GRASP) experiment by fusing RIG-5 and ZIG-8 to split GFP fragments. The resulting *spgfp11::rig-5* and *spgfp1-10::zig-8* double knock-in strain exhibited robust GFP fluorescence that precisely overlapped with ACR-16-AID-Scarlet neuronal puncta, thereby indicating the proximity of RIG-5 and ZIG-8 in the synaptic cleft **(Figure 2B)**.

Secondly, we assessed the impact of *rig-5* deletion on GFP-ZIG-8 localization. To completely abolish the *rig-5* gene activity, we engineered a full deletion mutant using CRISPR/Cas9 **(Supplementary Figure 2A)**. In this *rig-5(0)* mutant, GFP-ZIG-8 failed to concentrate at synapses in the ventral nerve cord, instead exhibiting a diffuse distribution along the neurites **(Figure 2C)**. Similarly, analyzing the distribution of a GFP-RIG-5 reporter in a *zig-8(0)* deletion allele revealed a dramatic loss of synaptic concentration **(Supplementary Figure 2B, Figure 2D)**. This underscores the essential roles of both RIG-5 and ZIG-8 in maintaining their synaptic localization.

Thirdly, we introduced point mutations in residues within the Ig1 domains previously demonstrated to disrupt the interaction between RIG-5 and ZIG-8^29^. We engineered a *gfp::rig-5(F75D)* allele and observed that both GFP-RIG-5(F75D) and Scarlet-ZIG-8 exhibited diffuse distributions along the neurites **(Figure 2E)**^29^. This suggested that disrupting the interaction between RIG-5 and ZIG-8 affects their synaptic localization. In a reciprocal experiment, we engineered a *Scarlet::zig-8(L77E)* allele, which also resulted in a diffuse distribution of both Scarlet-ZIG-8(L77E) and GFP-RIG-5 along neurites **(Figure 2E)**. Likewise, GFP-RIG-5(F75D) and Scarlet-ZIG-8(L77E) failed to concentrate at synapses in a *gfp::rig-5(F75D); Scarlet::zig-8(L77E)* double mutant **(Figure 2E)**. Taken together, these experiments strongly support the notion that the interaction between RIG-5 and ZIG-8 in the synaptic cleft is crucial for their synaptic localization.

### RIG-5 and ZIG-8 control ACR-16 clustering

Having identified RIG-5 and ZIG-8 in a genetic screen for mutants in ACR-16 localization, we aimed to precisely characterize their synaptic function **(Supplementary Figure 2A)**. In the *rig-5(0)* and *zig-8(0)* null mutants, as well as in the double *rig-5(0)*; *zig-8(0)* mutant, ACR-16-AID-Scarlet failed to cluster at neuron-neuron synapses **(Figure 3A)**. In contrast, the clustering of an ACR-16-AID-Scarlet reporter occurred normally at NMJ in *rig-5(0)* and *zig-8(0)* mutants **(Supplementary Figure 3A)**. This result is consistent with the known clustering mechanism of ACR-16 in muscle cells, which involves an intracellular scaffold composed of FRM-3/FARP and LIN-2/CASK^16^. To verify that the neuronal phenotype was independent of the Scarlet tag, we analyzed the *rig-5(0)* and *zig-8(0)* mutations using an ACR-16-AID-GFP knock-in reporter, and found similar clustering defects at neuronal synapses **(Supplementary Figure 3B)**. Additionally, given that the RIG-5(F75D) and ZIG-8(L77E) variants lose the ability to interact *in vivo* and accumulate at synapses, we investigated the impact of these mutations on ACR-16-AID-Scarlet distribution. As expected, ACR-16-AID-Scarlet did not cluster at synapses in these missense mutants **(Figure 3B, C)**. We concluded that RIG-5 and ZIG-8 control the concentration of ACR-16 specifically at neuron-neuron synapses of the ventral nerve cord, but not at NMJs.

**Figure 3.**
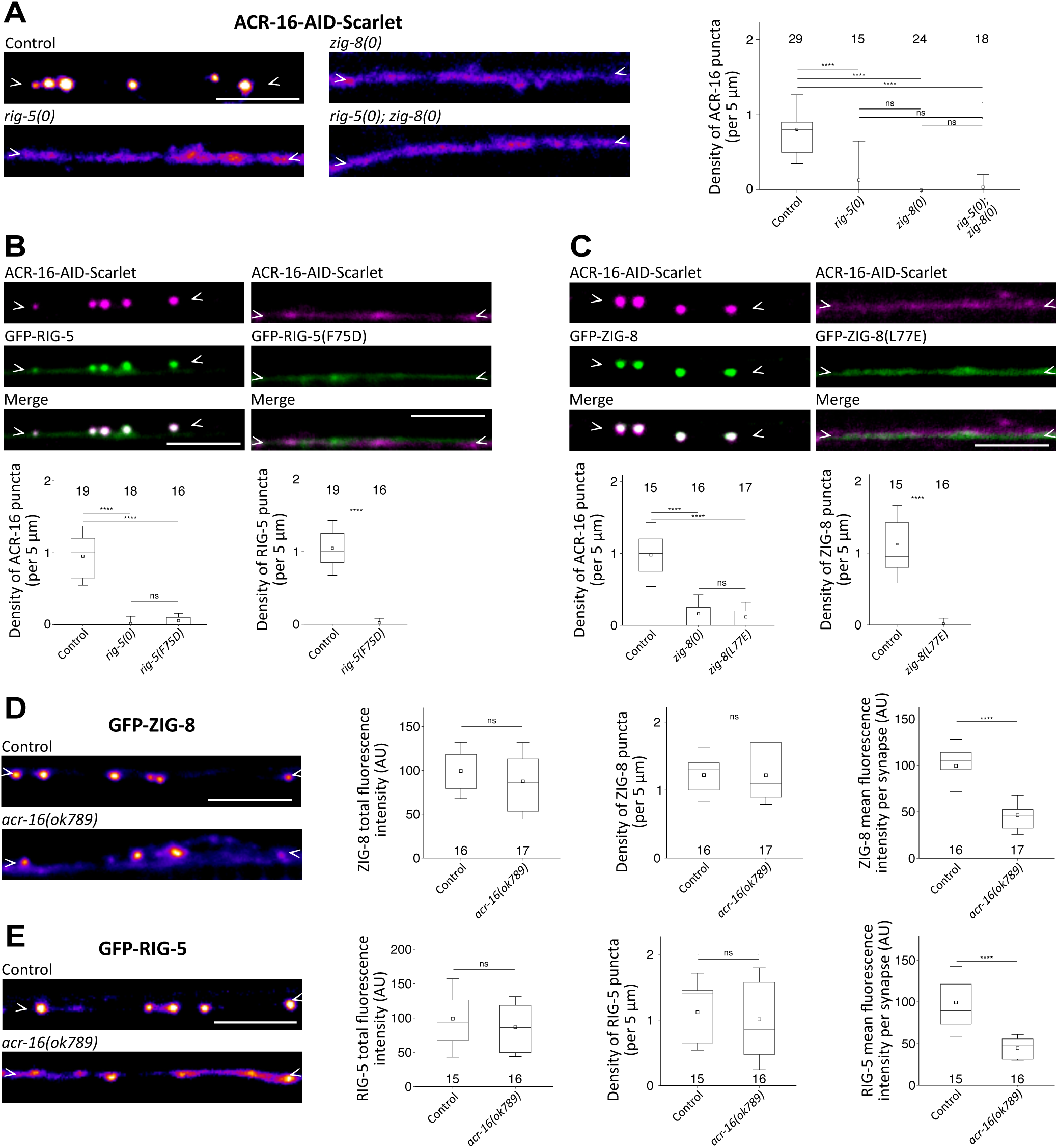
RIG-5 and ZIG-8 control ACR-16 clustering. **(A)** ACR-16-AID-Scarlet fluorescence shows clustering defects in neurons of the *rig-5(0)* and *zig-8(0)* mutants, as well as in the *rig-5(0); zig-8(0)* double mutant. The density of ACR-16 puncta is shown. **(B)** Clusters of ACR-16-AID-Scarlet and GFP-RIG-5 do not form when a mutation that prevents the RIG-5—ZIG-8 interaction is introduced (*gfp::rig-5(F75D)*). The density of ACR-16 and RIG-5 puncta is shown. **(C)** Clusters of ACR-16-AID-Scarlet and GFP-ZIG-8 do not form in the *gfp::zig-8(L77E)* mutant. The density of ACR-16 and ZIG-8 puncta is shown. **(D-E)** The GFP-ZIG-8 (D) and GFP-RIG-5 (F) fluorescence patterns were assessed in control and *acr-16(ok789)* mutant backgrounds. The total fluorescence intensity, puncta density and fluorescence intensity per synapse are shown. In this figure, an auxin treatment was applied to degrade ACR-16 in muscle cells. Data are presented as boxplots showing lower and upper quartiles (box), mean (square), median (center line) and standard deviation (whiskers); the number of worms is indicated for each condition; Kruskal-Wallis and Dunn’s test (A, B.1, C.1), Wilcoxon Mann-Whitney test (B.2, C.2, D.1, E.3), Student’s t-test (D.2,3, E.1,2); ns: non-significant, ****<0.00005. Scale bars: 5 µm.

Conversely, we investigated whether RIG-5 and ZIG-8 could concentrate at synapses in the absence of *acr-16*. We found that GFP-RIG-5 and GFP-ZIG-8 formed dimmer puncta at synapses, but also displayed a diffuse pattern between synapses in *acr-16* mutants **(Figure 3D, E)**. This suggests that ACR-16 is not critical for recruiting RIG-5 and ZIG-8 to synapses but may play a role in stabilizing their complex.

Synaptic adhesion molecules function during development and at mature synapses. To investigate whether RIG-5 and ZIG-8 have a developmental role, we analyzed larval stage 1 (L1) *C. elegans*. In newly born L1 larvae, BFP-RIG-5, GFP-ZIG-8 and ACR-16-AID-Scarlet highly colocalized in a punctate pattern, as in adult *C. elegans* **(Figure 2A, Supplementary Figure 4A)**. Moreover, we observed that *rig-5(0)* and *zig-8(0)* mutations prevented ACR-16-AID-Scarlet clustering **(Supplementary Figure 4B)**. Additionally, GFP-RIG-5 exhibited a diffuse pattern in the *zig-8(0)* mutant **(Supplementary Figure 4C)**. Similarly, GFP-ZIG-8 showed a diffuse pattern in the *rig-5(0)* mutant **(Supplementary Figure 4D)**. The consistent findings observed during both larval and adult stages suggest that RIG-5 and ZIG-8 control ACR-16 clustering throughout synapse development and maintenance.

The development of neuronal networks involves multiple coordinated steps, including neuronal specification, migration, axon growth and guidance, synaptic partner recognition, and synapse formation and maintenance. To explore the potential involvement of RIG-5 and ZIG-8 in the initial phases of neuronal network establishment, we assessed *rig-5(0)* and *zig-8(0)* single mutants for defects in neuronal specification, guidance and maintenance of the AVE, AVA and DB neurons. Our results revealed that these mutants did not exhibit noticeable defects in these processes **(Supplementary Figure 5A-E)**. These results are consistent with previous studies showing that RIG-5 and ZIG-8 contribute to nervous system development and maintenance, but in strict redundancy with other cell surface receptors and extracellular matrix components^34–36^. Taken together, our data strongly suggest that the main function of RIG-5 and ZIG-8 is to cluster the nicotinic receptor ACR-16 at specific cholinergic synapses.

### RIG-5 is presynaptic, and ZIG-8 postsynaptic

Our analysis has identified ACR-16 clusters in the AVA and DB neurons, notably at AVE to AVA synapses. This allowed us to investigate in which neurons *rig-5* and *zig-8* function. Using a neuronal gene expression atlas, we found that either *rig-5*, *zig-8*, or both are expressed in AVAs, DBs and most of their known synaptic partners **(Supplementary Figure 1C)**^19^. Specifically, *rig-5* was expressed in the AVE, AVD, DVA and AVA neurons, while *zig-8* was expressed in the AVE, AVA and DB neurons. The DB neurons exclusively express *zig-8*, suggesting a model in which RIG-5 might be presynaptic, and ZIG-8 might be postsynaptic at synapses onto AVAs and DBs.

To test this hypothesis, we performed a series of cell-specific rescue experiments, in single *rig-5(0)*, *zig-8(0)* or double *rig-5(0); zig-8(0)* mutant backgrounds, in which ACR-16 does not cluster properly. Firstly, we expressed BFP-RIG-5 in the AVE neurons of a *rig-5(0)* mutant and found that BFP-RIG-5 had a punctate distribution typical of AVE to AVA synapses, forming more than 20 puncta in the first quarter of the ventral nerve cord **(Figure 4A, D**, **Figure 1C, F)**. These BFP-RIG-5 puncta robustly colocalized with ACR-16-AID-Scarlet, suggesting that presynaptic expression of RIG-5 could rescue postsynaptic ACR-16 localization. Secondly, we introduced BFP-ZIG-8 in the AVAs of a *zig-8(0)* mutant. We observed an average of 80 puncta containing both BFP-ZIG-8 and ACR-16-AID-Scarlet along the ventral nerve cord, which likely correspond to synapses where ACR-16 clusters in AVA neurons **(Figure 4B, D**, **Figure 1C, F)**. This experiment suggested that expression of ZIG-8 in the postsynaptic AVA neuron could promote ACR-16-AID-Scarlet clustering. Thirdly, we co-expressed BFP-RIG-5 in the AVE neurons and GFP-ZIG-8 in the AVA neurons in the double *rig-5(0); zig-8(0)* mutant. We observed a sharp punctate distribution of BFP-RIG-5, GFP-ZIG-8, and ACR-16-AID-Scarlet, consistent with the synaptic pattern observed at AVE to AVA connections **(Figure 4C, D**, **Figure 1C, F)**.

**Figure 4.**
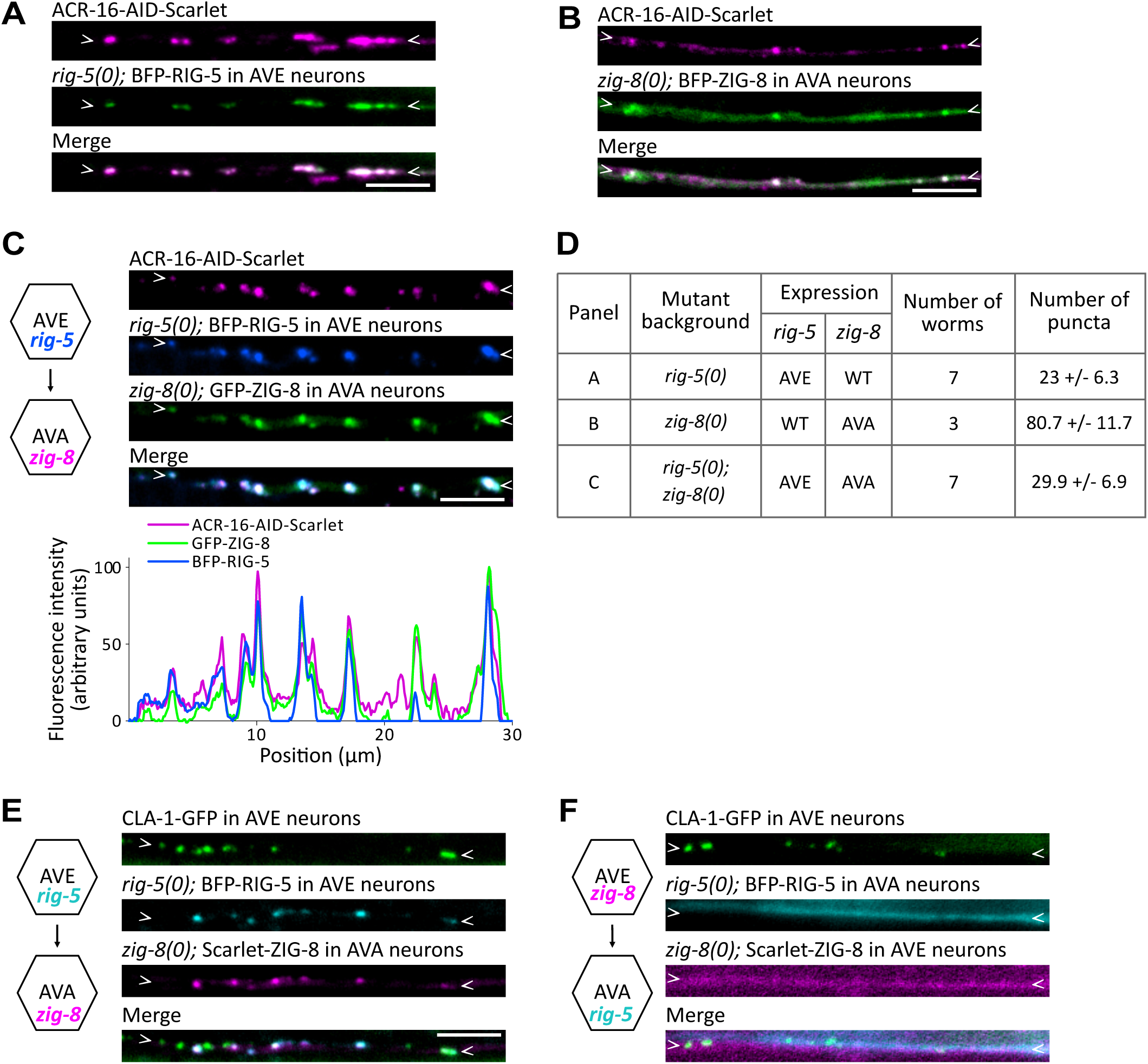
RIG-5 functions in presynaptic neurons and ZIG-8 in postsynaptic neurons. **(A)** Rescue of ACR-16-AID-Scarlet clustering in *rig-5(0)* mutants by expressing BFP-RIG-5 in AVE neurons. **(B)** Rescue of ACR-16-AID-Scarlet clustering in *zig-8(0)* mutants by expressing BFP-ZIG-8 in AVA neurons. **(C)** Rescue of ACR-16-AID-Scarlet clustering in *rig-5(0); zig-8(0)* double mutants by expressing both BFP-RIG-5 in AVE neurons and GFP-ZIG-8 in AVA neurons. The fluorescence intensity profile is shown below. Left: diagram representing cell-specific expression. **(D)** Number of ACR-16-AID-Scarlet clusters observed in the rescue strains (shown in A-C) along the ventral nerve cord. Data are shown as mean+/-SD. **(E)** Clusters of BFP-RIG-5 and Scarlet-ZIG-8 face an AVE-specific active zone marker (CLA-1-GFP), in *rig-5(0); zig-8(0)* double mutants expressing both BFP-RIG-5 in AVE neurons and Scarlet-ZIG-8 in AVA neurons. **(F)** In *rig-5(0); zig-8(0)* double mutants expressing both BFP-RIG-5 in AVA neurons and Scarlet-ZIG-8 in AVE neurons, no clusters are formed despite the presence of the AVE-specific CLA-1-GFP reporter. In panels A-C, an auxin treatment was applied to degrade ACR-16 in muscle cells. Scale bars: 5 µm.

To confirm the synaptic location of these puncta, we expressed BFP-RIG-5 in the AVEs along with an AVE-specific presynaptic marker, and Scarlet-ZIG-8 in the AVA, in the double *rig-5(0); zig-8(0)* mutant. We observed that BFP-RIG-5 and Scarlet-ZIG-8 formed puncta juxtaposed to the AVE presynaptic marker **(Figure 4E)**. To probe whether the orientation of the RIG-5—ZIG-8 trans-synaptic complex is important, we reversed their expression, introducing Scarlet-ZIG-8 in the presynaptic AVE neuron, and BFP-RIG-5 in the postsynaptic AVA neuron. Strikingly, we did not observe puncta of BFP-RIG-5 or Scarlet-ZIG-8 in this genetic background **(Figure 4F)**. This finding underscores that RIG-5 must be expressed in the presynaptic neuron and ZIG-8 in the postsynaptic neuron for correct localization to their respective pre- or postsynaptic membranes and efficient interaction at synapses. Overall, this set of cell-specific rescue experiments indicates that RIG-5 is presynaptic, and ZIG-8 postsynaptic, at least at AVE to AVA synapses.

### The Ig2 domain of ZIG-8 binds the ACR-16 extracellular domain

RIG-5 and ZIG-8 are adhesion proteins composed of 3 and 2 Ig-like domains, respectively. We analyzed the function of truncated RIG-5 and ZIG-8 proteins generated by CRISPR/Cas9 gene genome engineering, except for deletions of their Ig1 domains, which prevented their expression and were therefore not studied. Since ZIG-8 is required at the postsynaptic membrane, we investigated the role of its Ig2 domain in ACR-16-AID-Scarlet localization. Strikingly, in the absence of its Ig2 domain (*gfp::zig-8(Δ1Ig2)*), ZIG-8Δ1Ig2 formed synaptic clusters that overlapped with BFP-RIG-5, yet ACR-16-AID-Scarlet remained diffuse along the neurites **(Figure 5A)**. To exclude that this could be caused by the shortening of the protein, we replaced the ZIG-8 Ig2 domain with the human CD4 Ig2 domain (*gfp::zig-8(Δ1Ig2)::CD4(Ig2)*) and observed that ACR-16-AID-Scarlet also failed to cluster properly **(Figure 5A)**. These results indicate that the Ig2 domain of ZIG-8 is critical for recruiting ACR-16. In contrast, fragments of RIG-5 deleted for the Ig2, Ig3, or both Ig2 and Ig3 domains, were detected at synapses and effectively promoted ACR-16-AID-Scarlet concentration in the presence of full-length ZIG-8 **(Supplementary Figure 6A-C)**.

**Figure 5.**
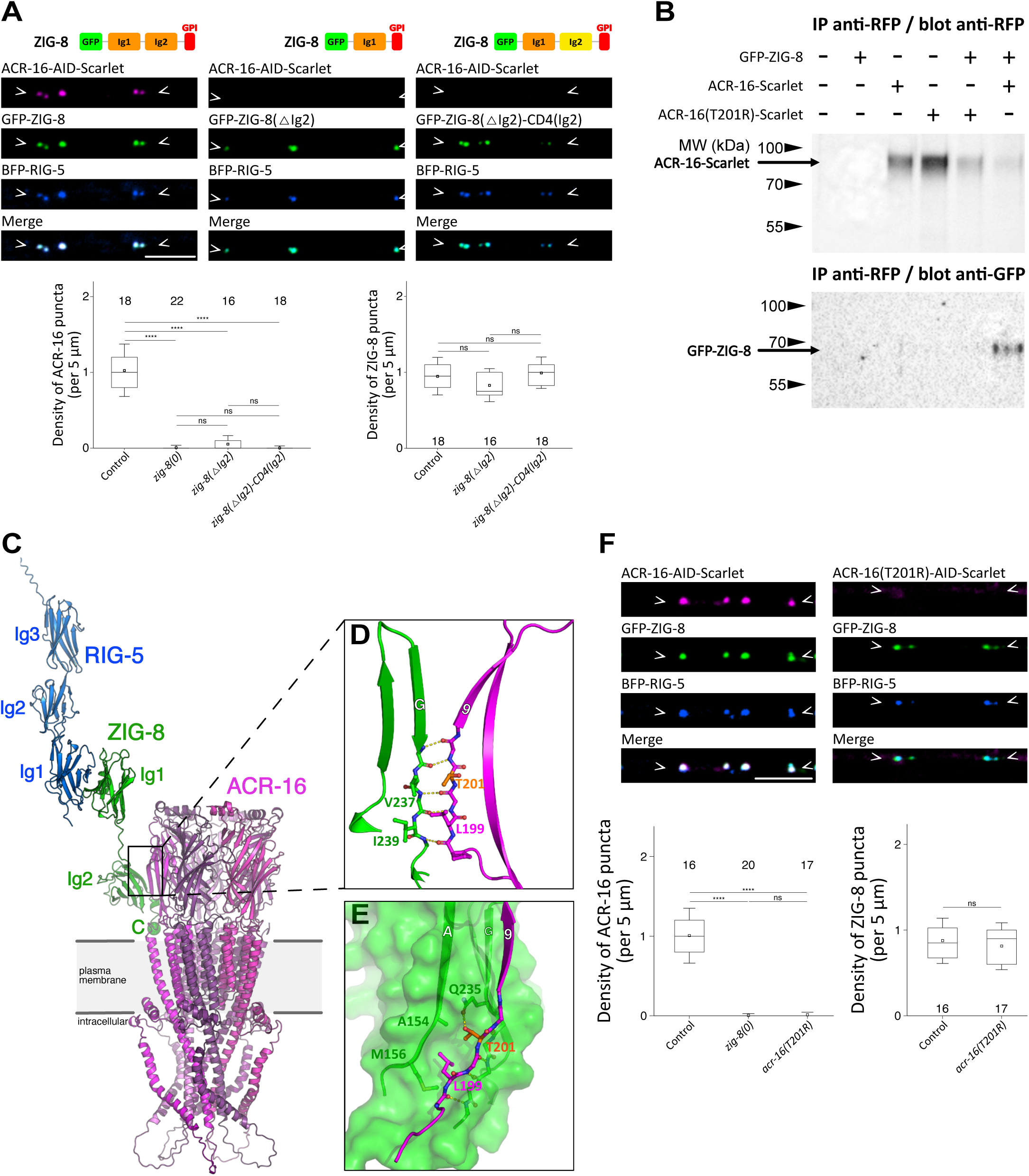
A direct interaction between the ZIG-8 Ig2 domain and the ACR-16 receptor. **(A)** ACR-16-AID-Scarlet receptors cannot form clusters when the Ig2 domain of ZIG-8 is deleted (*gfp::zig-8(Δ1Ig2)*) or when it is swapped with the Ig2 domain of human CD4 (*gfp::zig-8(Δ1Ig2)::CD4(Ig2)*). A schematic of ZIG-8 functional domains is shown for each knock-in condition. The density of ACR-16 and ZIG-8 puncta is shown. **(B)** Co-immunoprecipitation assay from HEK293 cells transfected or co-transfected with GFP-ZIG-8, ACR-16-Scarlet, or ACR-16(T201R)-Scarlet as indicated. The western blot below shows that GFP-ZIG-8 co-immunoprecipitates with ACR-16-Scarlet, but not with the mutated ACR-16(T201R)-Scarlet version. ACR-16-Scarlet and ACR-16(T201R)-Scarlet were co-expressed with the RIC-3 chaperone. **(C)** Cartoon representation of the composite RIG-5—ACR-16—ZIG-8 complex model as predicted by AlphaFold-multimer runs. ACR-16 protomers are drawn in various shades of magenta, while ZIG-8 is green and RIG-5 is blue. The C-terminal putative GPI anchor site of ZIG-8, labelled with a green sphere and letter “C”, is placed near the expected plasma membrane by Alphafold **(D)** ACR-16 strand 9 and the G strand of ZIG-8 Ig2 domain make an anti-parallel beta sheet. Dashed yellow lines represent H-bonds. **(E)**. Side chains of L199 and T201 in ACR-16 strand 9 sit in a complementary cavity on the ZIG-8 surface created by the A and G chains. Position of the T201 side chain is further stabilized by a hydrogen bond to the side chain of ZIG-8 Q235 (G strand). **(F)** The T201R mutation in ACR-16 (*acr-16(T201R)::aid::Scarlet*) prevents its clustering, while BFP-RIG-5 and GFP-ZIG-8 formed wild-type puncta. The density of ACR-16 and ZIG-8 puncta was assessed. In this figure, an auxin treatment was applied to degrade ACR-16 in muscle cells. Data are presented as boxplots showing lower and upper quartiles (box), mean (square), median (center line) and standard deviation (whiskers); the number of worms is indicated for each condition; Kruskal-Wallis and Dunn’s test (A.1, F.1), ANOVA and Tukey-hsd test (A.2, F.2); ns: non-significant, ****<0.00005. Scale bars: 5 µm.

Our results were compatible with a direct interaction between ZIG-8 and ACR-16. To test this hypothesis, we performed a co-immunoprecipitation experiment in HEK cells expressing both proteins. To prevent ZIG-8 from being released into the culture medium via cleavage of the GPI anchor, we replaced the C-terminal sequence of ZIG-8 containing the GPI anchor site with the transmembrane domain of human CD4 (GFP-ZIG-8-CD4(TM)). In addition, we coexpressed the RIC-3 chaperone to ensure proper intracellular trafficking of ACR-16-Scarlet^37^. The pull down of ACR-16-Scarlet efficiently co-immunoprecipitated GFP-ZIG-8-CD4(TM), hence supporting the existence of a physical interaction between ZIG-8 and ACR-16 **(Figure 5B)**.

To gain deeper insight into the interaction between the ZIG-8 Ig2 domain and ACR-16, we ran a structure prediction analysis using AlphaFold-Multimer. ACR-16 adopted a typical AChR structure, forming a pentamer composed of five subunits arranged around a central pore **(Figure 5C and Supplementary Figure 7A)**^38^. Interestingly, this analysis indicated potential interactions between ZIG-8 and each individual ACR-16 subunit at the base of the large N-terminal extracellular domain. The predicted interface involves several unconventional side-to-side, strand-to-strand hydrogen bond interactions, with the beta-strand G of ZIG-8 interacting with the beta-strand 9 of ACR-16 **(Figure 5D)**. This coupling is likely further stabilized by additional hydrogen bond and hydrophobic interactions involving atoms in the main and side chains of surrounding residues. For example, we observed that T201 and L199 side chains of ACR-16 sit in a mostly hydrophobic cavity formed between the A and G strands of ZIG-8’s Ig2 domain **(Figure 5E)**. Based on this, we predict that a bulkier side chain at these positions, such as an arginine at T201, would break this interaction.

To validate this predicted ZIG-8—ACR-16 interaction interface, we introduced a T201R point mutation into *acr-16::aid::Scarlet* (*acr-16(T201R)::aid::Scarlet*). To test whether ACR-16(T201R) was still functional, we used a loss-of-function mutation in *unc-29*, which encodes an essential subunit of the heteromeric L-AChRs present at the NMJ. When homomeric and heteromeric AChRs were non-functional in an *unc-29; acr-16* double mutant, animals were paralyzed, whereas the *unc-29* single mutant exhibited mild locomotion defects and the *acr-16* single mutant displayed normal locomotion **(Supplementary Figure 8)**. Importantly, the *acr-16(T201R)* mutant showed normal locomotion and did not exacerbate locomotion defects when combined with the *unc-29* mutation. These results indicated that the *acr-16(T201R)* mutation did not impair ACR-16 expression and function. To probe the interaction of the ACR-16(T201R)-AID-Scarlet variant with ZIG-8, we examined its clustering at neuron-neuron synapses *in vivo*. We found that the ACR-16(T201R)-AID-Scarlet variant failed to properly cluster at synapses labeled with BFP-RIG-5 and GFP-ZIG-8 **(Figure 5F)**. Consistently, the ACR-16(T201R)-AID-Scarlet receptor failed to co-immunoprecipitate GFP-ZIG-8-CD4(TM) in the HEK cell assay **(Figure 5B)**. Our data strongly suggest that the T201R mutation in *acr-16* disrupts its ability to interact with ZIG-8 *in vivo* at synapses and in a heterologous expression system.

To investigate whether ZIG-8 can directly cluster ACR-16 *in vivo*, we took advantage of the fact that ACR-16 is also expressed in muscle cells, but does not rely on RIG-5 and ZIG-8 for its synaptic localization. Rather, it involves an interaction of the large intracellular loop of ACR-16 with an intracellular scaffold comprising the FRM-3 multimodular protein. In *frm-3* null mutants, ACR-16-Scarlet is still present at the plasma membrane of muscle cells but fails to concentrate at NMJs and is not detected by fluorescence microscopy **(Figure 6B)**^16^. To test whether ZIG-8 is sufficient to cluster ACR-16, we engineered a mNeonGreen(mNG)-tagged chimeric protein between the ectodomain of the adhesion molecule NLG-1/neuroligin, which we previously demonstrated to localize to GABAergic NMJs of *C. elegans*, and the Ig2 domain of ZIG-8 with its GPI anchor signal **(Figure 6A, C)**^17^. Strikingly, expression of the mNG-NLG-1-ZIG-8(Ig2) chimera in the muscle cells of *frm-3* mutants caused the clustering of ACR-16-Scarlet at GABAergic NMJs. To confirm that the formation of these clusters did involve the ZIG-8—ACR-16 extracellular interface, we performed two control experiments. First, we introduced the T201R mutation in ACR-16-Scarlet and observed that mNG-NLG-1-ZIG-8(Ig2) no longer recruited ACR-16 at GABAergic NMJs **(Figure 6D, F-H)**. Second, we replaced the Ig2 domain of ZIG-8 by its Ig1 domain in the chimera and, although the mNG-NLG-1-ZIG-8(Ig1) chimera still localized to GABAergic NMJs, it failed to cluster ACR-16 **(Figure 6E, F-H)**.

**Figure 6.**
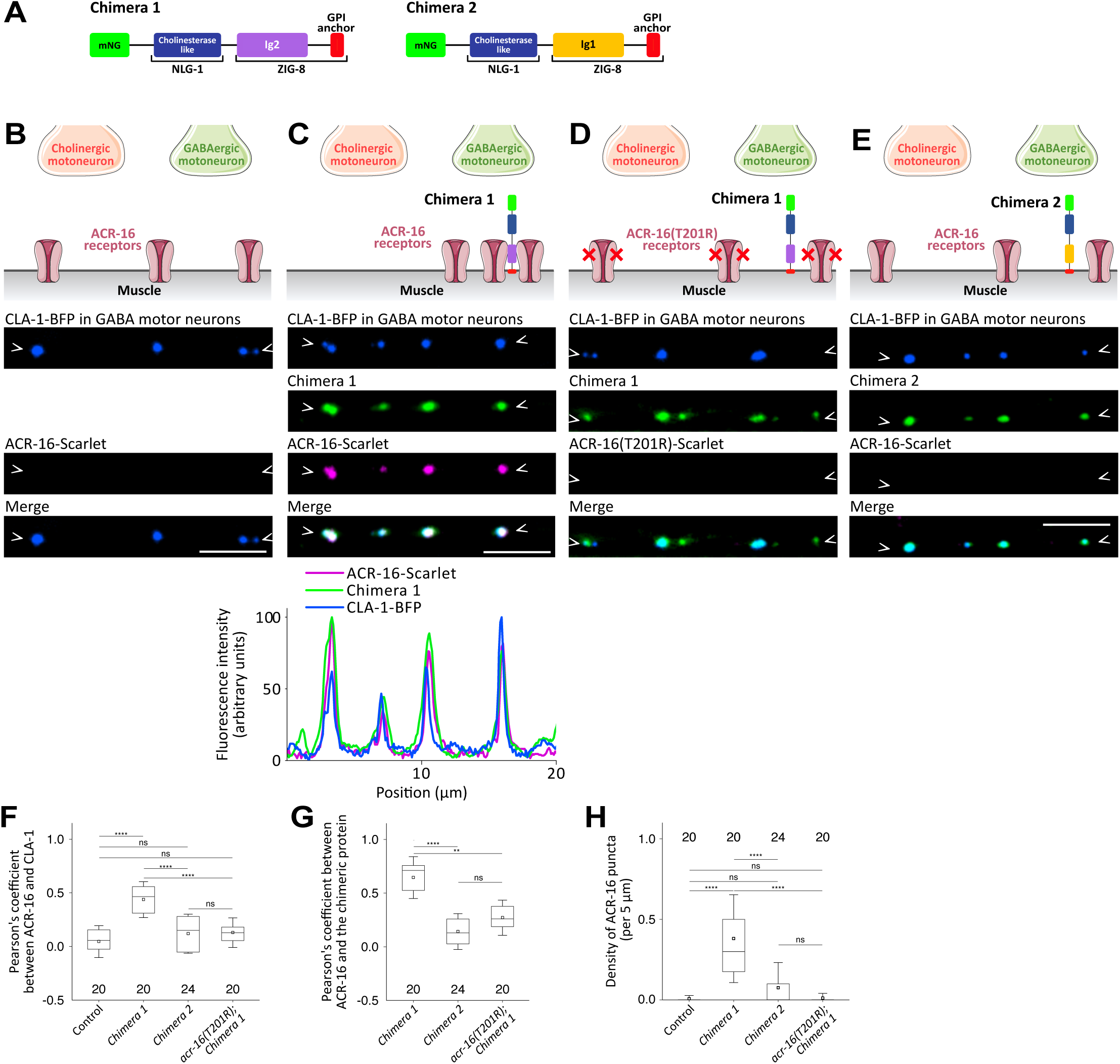
The ZIG-8 Ig2 domain is sufficient for ectopic clustering of the ACR-16 receptor at GABAergic neuromuscular junctions. **(A)** Domain composition of the two chimeric proteins: mNeonGreen (mNG), cholinesterase-like domain of NLG-1/neuroligin, Ig2 or Ig1 domain, and GPI anchor of ZIG-8. **(B)** ACR-16-Scarlet fails to cluster at NMJs in the *frm-3(0)* mutant, in the dorsal nerve cord. BFP-CLA-1 is expressed in GABA motor neurons to label GABAergic NMJs. **(C)** Chimera 1 colocalizes with ACR-16-Scarlet at GABAergic NMJs. The fluorescence intensity profile is shown below. **(D)** Chimera 1 localizes to GABAergic NMJs without ACR-16(T201R)-Scarlet. **(E)** Chimera 2 localizes to GABAergic NMJs without ACR-16-Scarlet. **(F-G)** Pearson’s coefficients indicate the degree of colocalization (ranging from -1: anti-correlation, to 1: complete colocalization) between ACR-16 and the BFP-CLA-1 presynaptic reporter (F) and between ACR-16 and the chimeric proteins (G). **(H)** The density of ACR-16 puncta is shown. Data are presented as boxplots showing lower and upper quartiles (box), mean (square), median (center line) and standard deviation (whiskers); the number of worms is indicated for each condition; Kruskal-Wallis and Dunn’s test (F-H); ns: non-significant, **<0.005, ****<0.00005. Scale bars: 5µm.

Taken together, these results support a direct mechanism whereby the Ig2 domain of ZIG-8 binds the base of the ACR-16 extracellular domain *in cis* on the postsynaptic membrane to cluster ACR-16 at neuron to neuron synapses.

## DISCUSSION

By focusing on cholinergic neuronal synapses in *C. elegans*, we have uncovered a novel paradigm for synaptic organization. Our unbiased genetic screen identified mutations in the *rig-5* and *zig-8* genes that cause dramatic defects in ACR-16 clustering. Our data indicate that RIG-5 and ZIG-8, the orthologs of the IgLONs cell adhesion molecules, form a trans-synaptic bridge within the synaptic cleft *in vivo* via their Ig1 domains. RIG-5 is tethered to the presynaptic membrane, whereas ZIG-8 is bound to the postsynaptic membrane. Remarkably, the Ig2 domain of ZIG-8 directly interacts with the extracellular domain of ACR-16 to localize this AChR at synapses. Altogether, our results uncover a novel mechanism of molecular assembly wherein a pair of IgLON cell adhesion molecules interact across synaptic membranes to directly cluster a postsynaptic cholinergic receptor.

### Clustering one cholinergic receptor by two distinct mechanisms

Acetylcholine is the most widely used neurotransmitter in *C. elegans*. Accordingly, the acetylcholine-gated ionotropic receptor family has been expanded and potentially contains around 30 AChR subunits that can assemble into homo- and heteromeric receptors^39^. By comparison, the human genome encodes only 16 AChR subunits, despite the many orders of magnitude greater complexity of the human brain. Only a few subunits have been assigned to specific synapses and characterized *in vivo* in *C. elegans*. The clustering of ACR-16 at the NMJ has been extensively studied. We have previously shown that it relies on an interaction with a synaptic intracellular scaffold comprising FRM-3, a FERM domain-containing protein orthologous to mammalian FARP1/2, and LIN-2, a MAGUK orthologous to CASK^16,18^. This intracellular scaffolding mechanism is strikingly different from the direct extracellular interaction observed with the ZIG-8 Ig2 domain at neuron to neuron synapses. Thus our results demonstrate that distinct mechanisms have evolved within the same species to control the clustering of a single ionotropic receptor type at different synapses.

### Limited characterization of neuronal AChR positioning mechanisms in mammals

In mammals, ionotropic nicotinic AChRs are expressed in a wide variety of neuronal and non-neuronal cell types, regulating various biological functions across tissues and systems^40^. In the central nervous system, they play important roles in controlling neural functions, including neuronal network activity and cognitive processing^41^. AChR are mainly present at presynaptic sites in axonal terminals, where they generally increase the release of many different neurotransmitters^42,43^. They are also found in postsynaptic regions and at extrasynaptic sites, where they are likely activated by volume transmission.

The mechanisms underlying the formation and positioning of neuronal AChRs within specific membrane microdomains remain unclear. Some studies have suggested that AChRs can be selectively clustered by classical postsynaptic intracellular scaffolds, such as those of the PSD95 family. However, it is still unknown whether this association is direct or indirect^44–46^. In addition, the PICK1 scaffolding protein has been shown to interact with the α7 cytoplasmic loop through an unconventional PDZ recognition mode, and to inhibit α7 surface expression in hippocampal GABAergic neuronal cultures^47,48^. In addition, PDZ and LIM domain 5 (PDLIM5) binds α7 AChRs via its PDZ domain, increasing their synaptic concentration and enhancing cholinergic neurotransmission^49^.

In contrast, no extracellular scaffolds have been described for neuronal AChRs. Nevertheless, AChRs interact with the lynx prototoxins, non-venomous proteins evolutionarily related to snake neurotoxins that can bind to the extracellular N-terminal domain of AChRs^50^. These proteins share a distinct “three finger” toxin fold and can be membrane-bound via GPI anchoring or secreted^51^. Due to their distinct binding specificity to AChRs and unique expression patterns in the mammalian brain, immune system, and other tissues, Lynx prototoxins have been proposed to modulate the function of specific AChR subtypes in a regionally specific manner. However, they have not been shown to control AChR localization. In *C. elegans*, it might be interesting in the future to test whether ZIG-8 binding modulates the function of ACR-16 in a similar manner, although ZIG-8 does not interfere with the ACR-16 agonist binding site based on *in silico* prediction.

Importantly we have identified for the first time that AChRs can be clustered within the synaptic cleft by interactions with their extracellular domain. Similar interactions of AChRs with IgLONs or other Ig-domain-containing proteins may control their subcellular localization in mammals.

### Neuronal roles of IgLONs across species

*rig-5* and *zig-8* are the sole members of the IgLON family in *C. elegans*. Up to now, no phenotype had been described in single null mutants, but discrete phenotypes have been reported for the maintenance of specific axons, in combination with mutants affecting cell surface and extracellular matrix proteins^34–36^. In contrast, this family, known as DIPs and Dprs, has expanded in *Drosophila melanogaster*^27,30^. There are 9 DIP and 21 Dpr paralogs able to form selective homo- and heterodimers through their Ig1 domains with varying affinities, such as in mammalian IgLONs^27,52–54^. Both DIPs and Dprs are present throughout the *Drosophila* nervous system, and are dynamically expressed in unique combinations in larval motoneurons, sensory neurons and muscle tissues^55–59^. This intricate protein interaction network has been shown to provide selective adhesion within the developing and mature *Drosophila* neural network. Notably, in the olfactory system, cell-type specific DIP-Dpr interactions support axonal self-adhesion and the precise sorting of olfactory receptor neurons into distinct territories^59^. In the visual system, interactions between DIP-alpha, Dpr 6 and Dpr10 have been shown to promote cell survival, control layer-specific innervation and modulate synapse density^60^. In addition, DIP-beta controls synaptic partner selection by establishing preferences for synapse formation between specific neuron types^61^. In the neuromuscular system, the interaction between DIP-alpha and Dpr10 plays a pivotal role in controlling the terminal branching of motoneurons onto larval body wall muscles and adult leg muscles, following specific patterns of innervation at both stages of development^56,58^. The combinatorial expression of DIPs and Dprs in motoneurons and their target muscle further suggests their involvement in regulating neuromuscular connectivity patterns. However, whether DIPs and Dprs function as bona fide synapse organizers in *Drosophila*, and the precise molecular mechanisms underlying their function in synaptic specificity, remain unclear.

There are five IgLON paralogs in mammals: IGLON5, LSAMP (Limbic System Associated Membrane Protein), OPCML (Opioid binding Protein/Cell adhesion Molecule Like), NTM (NeuroTriMin) and NEGR1 (Neuronal Growth Regulator 1). Proteomics studies of the adult mouse brain indicate that IgLONs are expressed in neurons, with some being expressed at high levels, while others are also expressed in glia^62^. In addition, IgLONs exhibit specific, regionally restricted patterns of expression in different brain regions^63^. The early onset of IGLON expression during embryogenesis, which persists into adulthood, suggests diverse functional roles throughout life^64^. Consistent with their distinct expression patterns, knock-out mouse models targeting each of the five IgLON genes exhibit significant but partially overlapping brain dysfunctions^65^. IgLONs are also widely expressed in the human brain and have been associated to a wide spectrum of neurological, neurodevelopmental, neuropsychiatric and neurodegenerative disorders, often involving synaptic dysfunction. These disorders include autoimmune encephalitis, mental retardation, autism spectrum disorders, major depressive disorder, schizophrenia and Alzheimer’s disease^65–68^. Paradoxically, the role of IgLONs in mammals is poorly understood.

IgLONs are capable of multiple homomeric and heteromeric interactions with varying affinities between members of the family^28,31–33^. Based on their neuronal expression, ability to engage in trans-cellular interactions, and co-expression in some brain areas, it has been widely suggested that IgLONs could bridge neuronal membranes at synapses and act as synaptic organizers^28^. Accordingly, a few studies support a synaptic role for IgLONs. Notably, proteomic characterization of rodent synaptic clefts has detected all IgLONs at glutamatergic synapses, and IGLON5 at GABAergic synapses as well^69–71^. At the ultrastructural level, LSAMP, OPCML, and NEGR1 have been shown to accumulate at synapses in various developing and adult brain regions^72,73^. In addition, OPCML has been identified at hippocampal synapses and found to promote dendritic stability through ephrin-cofilin signaling and regulation of F-actin dynamics^74^. Finally, a limited number of reports suggest that IgLONs modulate synaptic density in hippocampal neuronal cultures upon overexpression^75,76^. Despite compelling evidence for a role of IgLONs at synapses, this field remains largely unexplored.

Our current data show for the first time a pivotal role for the trans-synaptic interaction of *C. elegans* IgLONs in synaptic assembly. In addition, these results demonstrate that IgLONs are able to localize neurotransmitter receptors at the synapse via an unprecedented mechanism involving a direct interaction between an Ig-fold domain of a synaptic adhesion molecule and an ionotropic receptor. This finding is particularly significant given that the Ig superfamily, a large family of single-pass and GPI-anchored cell adhesion molecules, is widely present at synapses and may similarly recruit postsynaptic receptors^77^.

### RIG-5 and ZIG-8 synaptic IgLONs coordinate synaptic connectivity and synaptic organization

The anatomical simplicity of the *C. elegans* nervous system, together with its known inter-individual reproducible connectivity, provides unique opportunities to analyze synapses *in vivo* at individual resolution. In this study, we have analyzed a subset of synapses formed on the AVA and DB neurons that contain the ACR-16 AChR. AVAs are a pair of “command interneurons” that integrate numerous sensory inputs, connect directly to motoneurons and regulate locomotion^20,78^. These neurons are among the most connected neurons in *C. elegans*, receiving more than 200 inputs, primarily from sensory neurons and other interneurons. However, our results have identified a specific subset of AVA synapses containing RIG-5, ZIG-8 and ACR-16. Overall, our data highlight a strong specificity in the molecular composition of synapses among AVA neurons, raising the question of how RIG-5 and ZIG-8 are trapped at specific synapses.

A first level of control relies on the specific expression of *rig-5* and *zig-8* in defined classes of neurons **(Supplementary Figure 1C)**. For example, *rig-5* is expressed in the presynaptic AVE neurons but not in the AVB neuron, a cholinergic interneuron that provides 20-30 synaptic inputs but does not use ACR-16 as a receptor at these synapses **(Supplementary Figure 1G)**. However, cell-specific expression does not explain how RIG-5 and ZIG-8 cluster in specific locations. These molecules are bound to membranes by a lipid anchor and are likely to diffuse freely along the plasma membranes. During development, transient contact between pre- and postsynaptic membranes may be sufficient to engage RIG-5 and ZIG-8 interactions in *trans*, and nucleate synaptic differentiation. Accordingly, no cluster of RIG-5 or ZIG-8 is observed when either molecule is absent **(Figure 2C, D)**.

Additionally, our data raise the question of how the concentration of RIG-5 and ZIG-8 is achieved. While synaptic scaffolding molecules have often been shown to multimerize in different stoichiometric complexes^5,7^, structural studies of *C. elegans* RIG-5 and ZIG-8, *Drosophila* DIPs and Dprs, and mammalian IGLONs suggest that they solely form dimers through their Ig1 domains, rather than higher-order multimers^28,29,31–33^. Nevertheless, our structural prediction is consistent with each molecule of ZIG-8 interacting *in trans* with RIG-5, and laterally *in cis* with each subunit of ACR-16, potentially forming a pentameric RIG-5—ZIG-8—ACR-16 assembly **(Figure 5C)**. This model suggests that ACR-16 could stabilize the whole complex through pentamerization. Accordingly, in the *acr-16* mutant, GFP-RIG-5 and GFP-ZIG-8 display reduced synaptic concentration and a diffuse pattern between synapses, suggesting stabilization defects **(Figure 3D, E)**.

Although our simple model of protein interactions between IgLONs and ACR-16 is sufficient to explain the formation of postsynaptic AChR clusters, the lack of any transmembrane and intracellular region is difficult to reconcile with coupling to presynaptic differentiation. A recent analysis of the nematode extracellular interactome identified several RIG-5 and ZIG-8 interacting partners, including potential presynaptic proteins^79^. It will be interesting to test these candidates and possibly identify new binding partners of RIG-5 and ZIG-8 to account for the coordinated differentiation of pre- and postsynaptic specializations.

In any case, our genetic strategy in *C. elegans* has uncovered novel functions for IgLON molecules and a novel paradigm for AChR clustering. Since all these molecules and protein domains have been conserved during evolution, it will be interesting to test to what extent these mechanisms are utilized in the mammalian brain.

## ACKNOWLEDGEMENTS

We thank members of the Bessereau lab for providing feedback, Mélissa Cizeron for advice on image analysis, Laure Granger for expert advice on cell culture and biochemistry, Driss Laabid for technical assistance, and Océane Romatif, Delphine Le Guern and Camille Vachon for strains. We are grateful to Elise Forgues, Cecile Chatras, Mathieu Ben Abu and Manon Courtieux for their help during their respective internships. We thank Pierre-Jean Corringer for critical reading of the manuscript, Mei Zhen and Jun Meng for providing information on neuronal promoters, Alexander Gottschalk for sharing plasmids and Eviatar Yemini for advice on NeuroPAL. We thank the Caenorhabditis Genetics Center (CGC), funded by NIH Office of Research Infrastructure Programs (P40 OD010440), for providing strains. We thank the SFR Biosciences (University Lyon 1 CNRS UAR 3444 INSERM US8, ENS de Lyon), Matthieu Caron and Francesca Palladino for access to equipment. We thank Le Centre d’Imagerie Quantitative Lyon-Est (LyMIC-CIQLE, Lyon, France) imaging facility for support and access to equipment, and Camilla Luccardini for technical assistance. We thank the SEGiCel for strains (SFR Santé Lyon Est CNRS UAR 3453, Lyon, France). This work was supported as follows: M.M.: fellowship from the French Ministry of Research; BPL: LABEX Cortex (ANR-11-LABX-0042) of University Lyon 1, within the program “Investissements d’Avenir” (ANR-11-IDEX-0007); EÖ: National Institutes of Health, National Institute of Neurological Disorders and Stroke grant R01 NS139060 grant; JLB: Équipe FRM 2023 EQU202303016267, ANR Synapunct ANR-22CE16-0024-01, ERC_Adg C.NAPSE #695295, ANR-11-LABX-0042/ANR-11-IDEX-0007.

## AUTHOR CONTRIBUTIONS

Conceptualization: M.M., B.P.-L. and J.-L.B.; Methodology: M.M., A.W., E.Ö., J.-L.B. and B.P.-L.; Investigation: M.M., L.P. and B.P.-L.; Formal analysis: M.M., L.P., E.Ö. and B.P.-L.; Visualization: M.M., E.Ö., J.-L.B. and B.P.-L.; Writing – Original Draft: M.M., J.-L.B. and B.P.-L.; Writing – Review & Editing: M.M., A.W., E.Ö., J.-L.B. and B.P.-L.; Funding Acquisition: E.Ö., J.-L.B. and B.P.-L.; Supervision: J.-L.B. and B.P.-L.

## DECLARATION OF INTERESTS

The authors declare no competing interests.

## DECLARATION OF GENERATIVE AI AND AI-ASSISTED TECHNOLOGIES IN THE WRITING PROCESS

During the preparation of this work the authors used ChatGPT - OpenAI and DeepL in order to improve the readability and language of the manuscript. After using these tools, the authors reviewed and edited the content as needed and take full responsibility for the content of the publication.

## MATERIALS AND METHODS

### Strains

All experiments were performed at 20°C. Strains were grown on nematode growth medium (NGM) agar plates seeded with *Escherichia coli* OP50^80^. The wild-type reference strain was *C. elegans* N2 Bristol. All strains and alleles used in this study are described in **Supplementary Table S1 and S2**, respectively.

### Strains generated by EMS mutagenesis

Several alleles of *rig-5* and *zig-8* were identified through genetic screening for mutants showing defects in ACR-16-AID-Scarlet clustering: *rig-5(kr573[Q16*])*, *rig-5(kr849[P191S])*, *zig-8(kr850[Q92*]), zig-8(kr851[A68T])* and *zig-8(kr852[I239F])*.

### Allele generation by CRISPR/Cas9 genome engineering

All knock-in alleles were generated according to a standard protocol^81^. crRNA was designed using Benchling software and synthesized by Integrated DNA Technologies (IDT). To form a triplex, a mix of 2.8 μL crRNA (34 μM), 5 μL tracrRNA (18 μM) and 0.5 μL Cas9 nuclease (10 μL/μg) (IDT) was incubated for 15 min at 37 °C. For deletions, two crRNA were used at both ends of the target sequence, with a triplex made of 1.4 μL of each crRNA. A mix containing 2.2 μL single-strand repair template (1 μg/μl) or 500 ng double-strand repair template, with 800 ng of pRF4 plasmid, and molecular biology grade water up to 20 μL was added to the triplex. This mix was injected into adult *C. elegans*. Worms with a roller phenotype in the next generation were isolated and tested by PCR. All gene edits were confirmed by sequencing. Finally, the edited strains were outcrossed once to remove nonspecific background mutations. A list of crRNA is provided in **Supplementary Table S3**.

### Plasmids

The plasmids constructed for this study are described in **Supplementary Table S4**. All constructs were verified by Sanger sequencing from the GATC company. pMM46,48,53,57 derive from pcDNA3.1. Plasmid sequences are available upon request.

### Tissue-specific expression

For tissue-specific expression, the promoters used were as follows:

- *Pmyo-3*: body wall muscle cells
- *Punc-17*: cholinergic motoneurons
- *Punc-129*: D-type motoneurons (DA and DB type)
_-_ *Ppept-3*: AVE neurons^82^

For specific expression in AVA and AVB neurons, the Cre-loxP recombination strategy was used, with promoter combination as follows: *Pflp-18* and *Pgpa-14* (AVA neurons), *Ptkw-40* and *Plgc-55* (AVB neurons)^24,83,84^. Promoter boundaries are described in **Supplementary Table S5**.

### Generation of single-copy insertion alleles

Single-copy insertion alleles were generated by the miniMos method^85^. Briefly, adult *C. elegans* were injected with a mix containing 15 ng/μL of the plasmid of interest (containing promoters and open reading frames fused to fluorescent proteins), 50 ng/μL pCFJ601 (Mos1 transposase), 10 ng/μL pMA122 (negative selection marker *Phsp16.2::peel-1*), and 2.5 ng/μL pCFJ90 (*Pmyo-2::mCherry*). Neomycin (G418) was added to plates 24 h after injection at 1.5 μg/μL final concentration. Candidate plates were heat-shocked for 2h at 34 °C. Worms with an insertion were isolated and homozygosed.

### Multicopy and extrachromosomal array lines

Extrachromosomal array alleles were generated by injecting a mix of plasmids of interest as depicted in the Supplementary Table S2.

For *acr-16* transcriptional reporter (*krIs83[Pacr-16::aid::mNG]*, **Supplementary Figure 1B**), a strain bearing a *Pacr-16::aid::mNG* extrachromosomal array was randomly integrated by X-ray irradiation (40 Gy).

### Auxin-induced cell specific degradation

For all images except in Fig. 4E, F, Fig. 6, Supp. Fig.1B, Supp. Fig. 3A, Supp. Fig. 4 and Supp. Fig. 5, worms were grown on auxin plates to degrade ACR-16-AID-Scarlet at NMJs. Specifically, adult *C. elegans* were transferred to auxin plates, and their progeny was analyzed. Auxin plates were prepared by adding auxin (indole-3-acetic acid, Sigma-Aldrich) from a 400 mM stock solution in ethanol to NGM, at the final concentration of 1 mM^21^.

### Neuron-type Specific Illumination (NeuroSIL)

ACR-16 was tagged with AID and Scarlet sequences, along with three copies of spGFP11 (ACR-16-AID-Scarlet-spGFP11). The spGFP1-10 moiety was expressed in the cytoplasm of AVA or DB neurons using selected combinations of promoters, as described above. All ACR-16 clusters displayed Scarlet labelling but only ACR-16 clusters formed in the AVAs or DBs, exhibited GFP fluorescence upon reconstruction. Two to four transgenic lines were observed, and/or used for quantification: Fig. 1E (2 lines), Fig. 1H (4 lines) and Supplementary Fig. 1F (2 lines).

### Microscopy imaging and quantification

For spinning disk imaging, young adult hermaphrodites were used, except in Supplementary Fig. 4, in which L1 larvae were imaged 1-2 h after hatching. Live worms were mounted on 2% agarose dry pads with 2% (for young adults) or 5% (for L1 larvae) polystyrene beads (Polybeads, Polysciences) in M9 buffer. Worms were observed using an Andor spinning disk system (Oxford Instruments) installed on a Nikon-IX86 microscope (Olympus) equipped with a 60×/NA 1.42 oil-immersion objective and an Evolve electron-multiplying charge-coupled device camera. Each animal was imaged with IQ software (APIS Informationstechnologien) or Metamorph software (version 7.10.5.476) as a stack of optical sections (0.2 μm apart) across the whole thickness of the ventral or dorsal nerve cords. All images were processed using Fiji software (v2.0) and correspond to the sums of all slices with signal along the Z-axis^86,87^.

Acquisition settings were the same across genotypes for quantitative analysis. Image quantification was performed using a macro in Fiji. For total fluorescence intensity measurements, a 50 μm (wide) × 3 μm (high) rectangle containing solely the nerve cord (first quarter) was cropped and projected as a sum in Z. The mean intensity was then projected in the x direction (orthogonal to the cord), and the intensity was measured as the area under the curve, after background exclusion^88^. Data are presented as a percentage of the average fluorescence relative to that of the wild type.

For puncta density and intensity quantification, a CORSEN analysis was used in Fig. 3D, E89. Due to the diffuse localization pattern observed in Fig. 2C-E, Fig. 3A-C, Fig. 5A, F, puncta density was quantified as follows: the cropped image (50 μm wide × 3 μm high) was converted to a one-dimensional axis along the y axis in Fiji (ImageJ). Plot profiles were generated for each image and saved as csv files. To determine puncta density, the csv files were computed by an R script to apply a threshold, define the background signal and count the number of peaks above this threshold. The threshold was the same across genotypes for each quantitative analysis. In Fig. 1 and 4, the total number of puncta along the ventral nerve cord was quantified manually. The quantification of BFP-tagged proteins could not be assessed automatically due to a low signal-to-noise ratio.

For colocalization analysis, images were captured along 50 μm of the dorsal nerve cord anterior to the vulva. Fluorescence intensity along the cord was evaluated with the Plot Profile Fiji plugin. For each channel, values along the x axis were normalized to the maximal intensity value. Data are presented as minimum to maximum values for animals of each genotype. Colocalization between two channels was analyzed using Pearson’s correlation coefficient as previously described^17^. For quantification, data are presented as box plots showing lower and upper quartiles (box), mean (square), median (center line) and standard deviation (whiskers). Statistical tests were performed using RStudio version 2022.12.0 (R version 4.0.5).

For rescue experiments, one to three lines transgenic lines were observed, and/or used for quantification: Fig. 4A (1 line), Fig. 4B (1 line), Fig. 4C (2 lines), Fig. 4E (3 lines) and Fig. 4F (2 lines).

The NeuroPAL (Neuronal Polychromatic Atlas of Landmarks) strain was used to identify *acr-16*-expressing neurons based on a *Pacr-16::aid::mNG* transcriptional reporter in head, body and tail regions, by comparison to the reference manual^22^.

### Transfection and Immunoprecipitation

HEK293 cells were maintained in DMEM medium (Invitrogen) supplemented with 10% inactive fetal bovine serum (Gibco) and 1% penicillin-streptomycin (Sigma, P0781) in a humidified atmosphere with 5% CO_2_ at 37 °C. The plasmids pMM46 (expressing RIC-3), pMM48 (expressing ACR-16-Scarlet), pMM53 (expressing GFP-ZIG-8 with the GPI anchor signal swapped to the transmembrane region of human CD4) and pMM57 (expressing ACR-16(T201R)-Scarlet) were transfected or co-transfected using a Calcium Phosphate transfection method. RIC-3 was systematically co-transfected with ACR-16. Note that the yield of ACR-16 and ZIG-8 expression was reduced upon their co-transfection (Fig. 5B). The cells were harvested at 48 hours after transfection and lysed with ice-cold WRB1 immunoprecipitation buffer (50 mM HEPES pH 7.7, 100 mM NaCl, 50 mM KCl, 1mM EDTA, one tablet of protease inhibitor cocktail per 50 mL of buffer) containing 2% Triton X-100 on ice for 2h, then centrifuged at 16,000 g for 15 min at 4 °C. The cell lysates were diluted with WRB1 in a 1:1 ratio. 20 μL of prewashed anti-RFP-Trap-A beads (Chromotek) were added and incubated overnight at 4 °C with gentle rotation. The beads were collected by centrifugation at 2,000 g for 2 min at 4 °C and washed 3 times with WRB1. The immunoprecipitated proteins were eluted in Laemmli buffer with 2-mercaptoethanol, and analyzed by Western blotting after boiling for 5 min at 95 °C. The primary anti-GFP monoclonal antibody (Roche, 1814460001) was used at a 1:2,000 dilution and the primary anti-RFP monoclonal antibody (Chromotek, 6G6) was used at a 1:1,000 dilution. Horseradish peroxidase-conjugated anti-mouse (K4000; DAKO) was used as secondary antibody at 1:3000 dilution. Images of blotting membranes were captured by Bio-rad ChemiDoc XRS + imaging system.

### Structure prediction and analysis

Protein complex structure predictions were performed with AlphaFold-Multimer, using the ColabFold implementation version 1.5.2 running Alphafold release 2.3.1^90–92^. For the prediction, we used the default settings, including the use of template information, relaxation of the predicted structures using amber force fields, and use of both paired and unpaired multiple sequence alignments. We ran alphafold predictions of the ACR-16 pentamer, and various ectodomain complexes of ACR-16 and ZIG-8, all returning models that were in close agreement. Alphafold reported high confidence values for the interfaces: the iPTM value for the ACR-16 pentamer, the 1:1 complex of ACR-16 and ZIG-8 and the ACR-16 pentamer with a ZIG-8 molecule were 0.849, 0.865 and 0.845. An example of the sequence coverage, pLDDT, and predicted alignment error (PAE) plots are shown in **Supplementary Figure 7B**. Analysis of the structures and creation of composite models (including the ZIG-8-RIG-5 crystal structure) were performed using the molecular graphics program PyMOL (v. 3.0.4)^29,93^.

### Locomotion test

Worm locomotion was assessed on plates by gently tapping the head and tail of each animal using a platinum wire mounted on a pick, as well as following mechanical tapping of the Petri dish. The assessments were conducted by three experimenters who were blinded to the genotype.

## SUPPLEMENTARY FIGURE LEGENDS

**Supplementary Figure 1.**
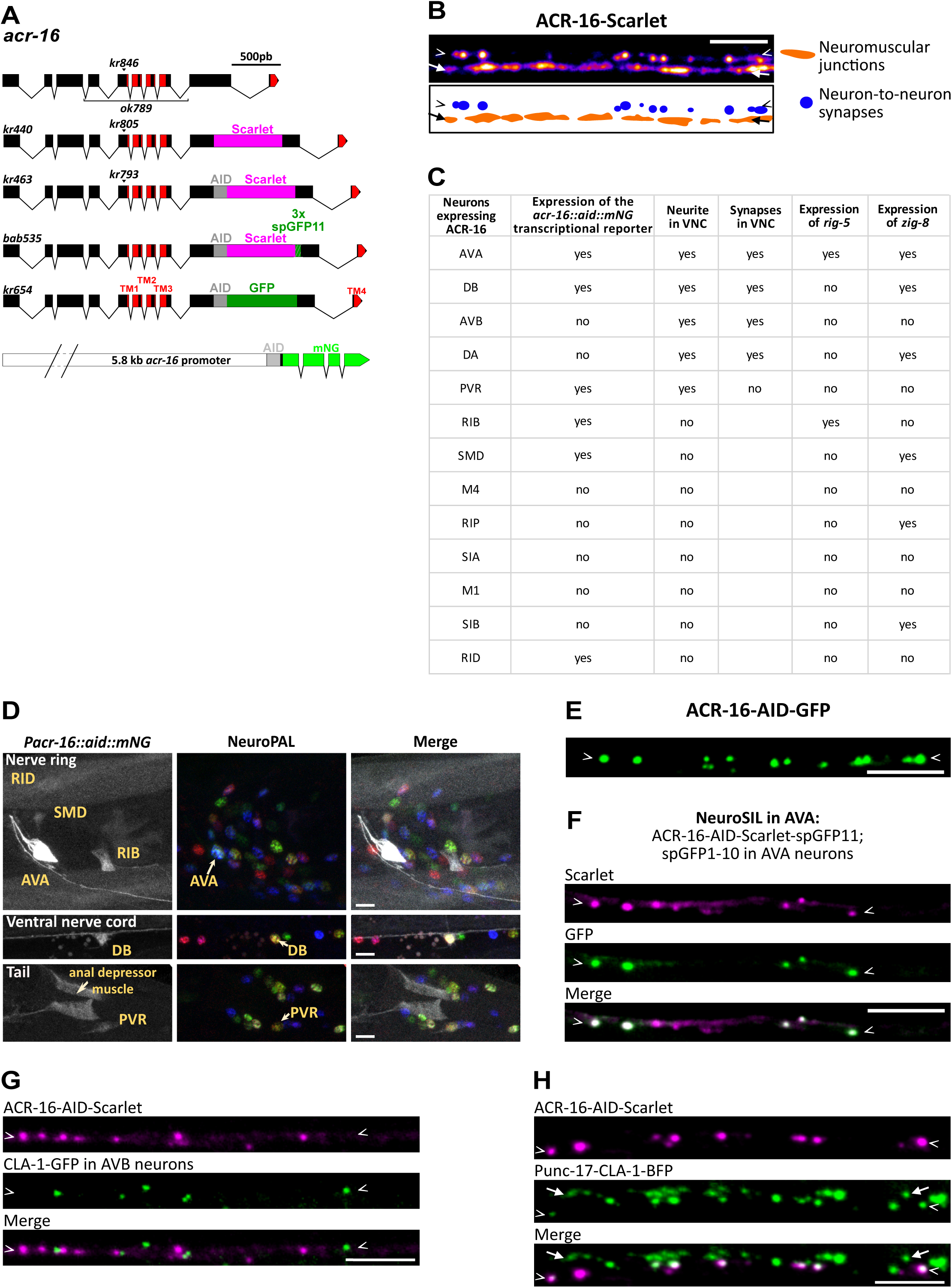
Analysis of ACR-16 expression and synaptic localization. **(A)** Genomic organization of the *acr-16* locus (boxes represent exons with sequences of the four transmembrane domains colored in red), and the positions of different tags and mutants used in this study (arrowheads represent point mutations; the bracket represents a deletion). Bottom: transcriptional reporter. Scale bar: 500 bp. **(B)** Spinning disk microscopy images and schematic of ACR-16-Scarlet fluorescence along the ventral cord at NMJs (orange dots, arrows) and neuron-neuron synapses (blue dots, arrowheads). **(C)** Table depicting neurons that display *acr-16* expression, whether they form neurites and synapses in the ventral nerve cord, and whether they express *rig-5* and *zig-8.* Expression data has been extracted from the CeNGEN atlas using a threshold of 2 (yes = detected with a threshold of 2, no = not detected). Connectivity data from White *et al.*, 1986. **(D)** The *acr-16* transcriptional reporter (*krIs83[Pacr-16::aid::mNG]*) was analyzed in the NeuroPAL background in the head, ventral nerve cord and tail, following auxin-induced degradation in muscle. **(E)** Punctate pattern of ACR-16-AID-GFP. **(F)** NeuroSIL images in AVA neurons showing Scarlet fluorescence and GFP reconstruction (ACR-16-AID-Scarlet-spGFP11, spGFP1-10 in AVAs). For (E-F), the puncta number was assessed in Figure 1C. **(G-H)** Endogenous ACR-16-AID-Scarlet and a CLA-1-GFP presynaptic reporter expressed in AVB neurons (G) and cholinergic neurons (H). Scale bars: 5 µm.

**Supplementary Figure 2.**
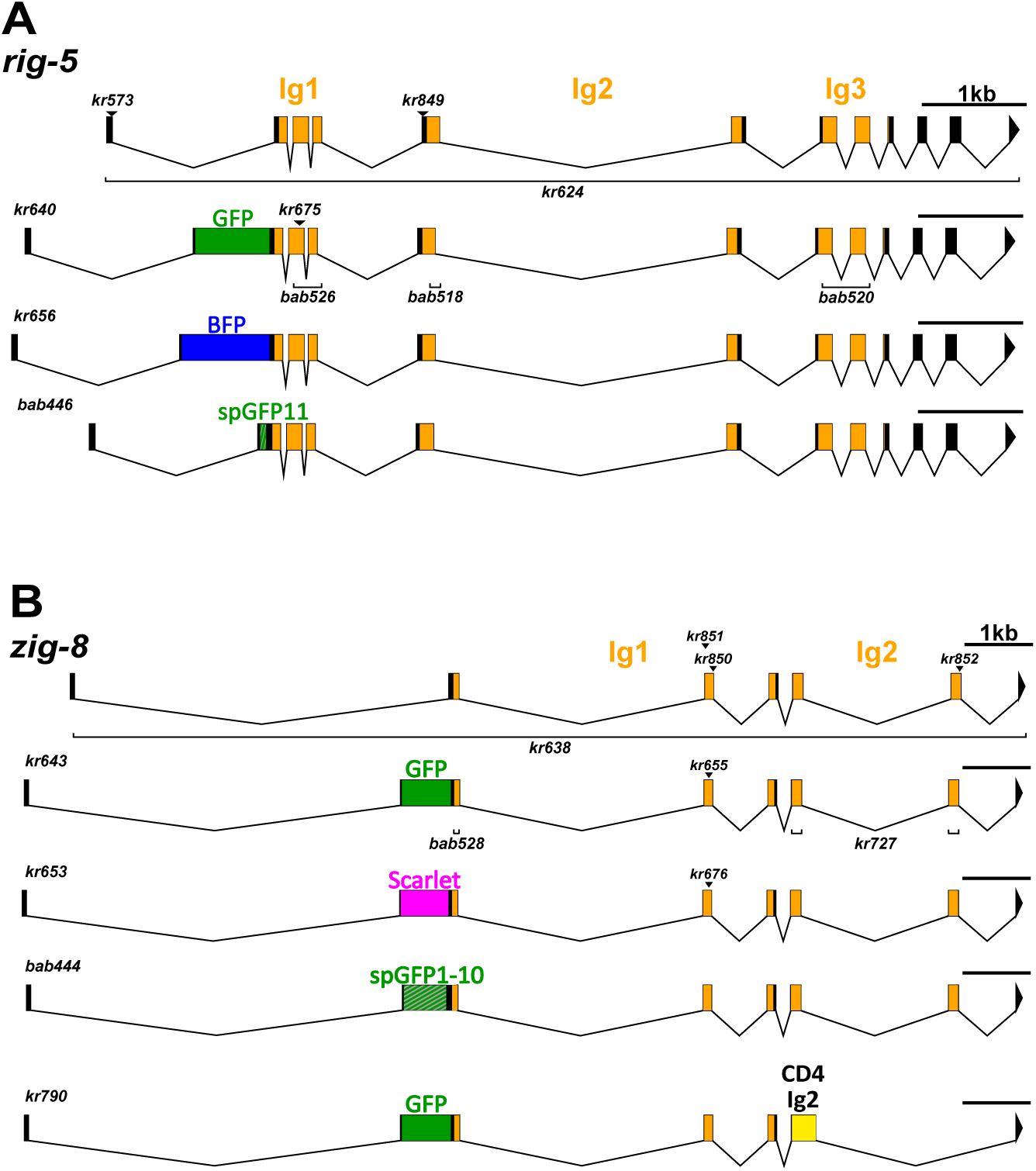
The *rig-5* and *zig-8* loci. **(A)** Genomic organization of *rig-5* and the positions of different tags and mutants used in this study. **(B)** *zig-8* locus, tags and mutants, including an insertion used in this study. Boxes represent exons with sequences of the Ig domains colored in orange. Arrowheads: point mutations; brackets: deletions. The Supplementary Table S2 contains a brief description of *rig-5* and *zig-8* alleles.

**Supplementary Figure 3.**
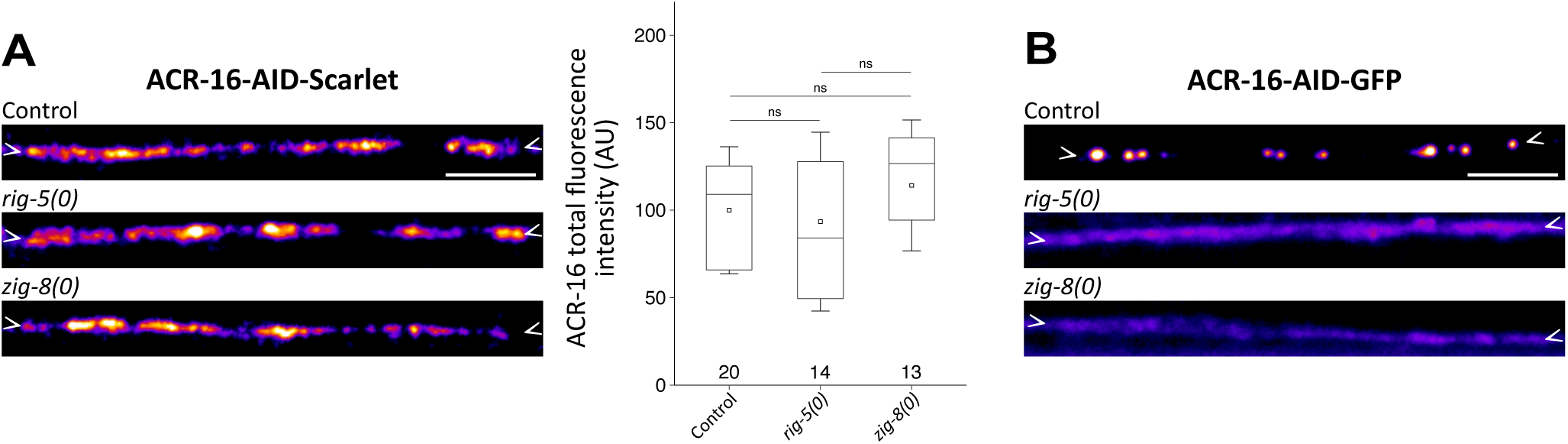
*rig-5* and *zig-8* mutations prevent ACR-16 clustering at neuron-neuron synapses but not at neuromuscular junctions. **(A)** ACR-16-AID-Scarlet labelling at neuromuscular synapses of the dorsal nerve cord, with quantification of the total fluorescence intensity in a control strain and *rig-5(0)* and *zig-8(0)* mutants. **(B)** ACR-16-AID-GFP fluorescence shows clustering defects in neurons of the *rig-5(0)* and *zig-8(0)* mutants following auxin-mediated degradation in muscle cells. Data are presented as boxplots showing lower and upper quartiles (box), mean (square), median (center line) and standard deviation (whiskers); the number of worms is indicated for each condition; Kruskal-Wallis and Dunn’s test (A); ns: non-significant. Scale bars: 5 µm.

**Supplementary Figure 4.**
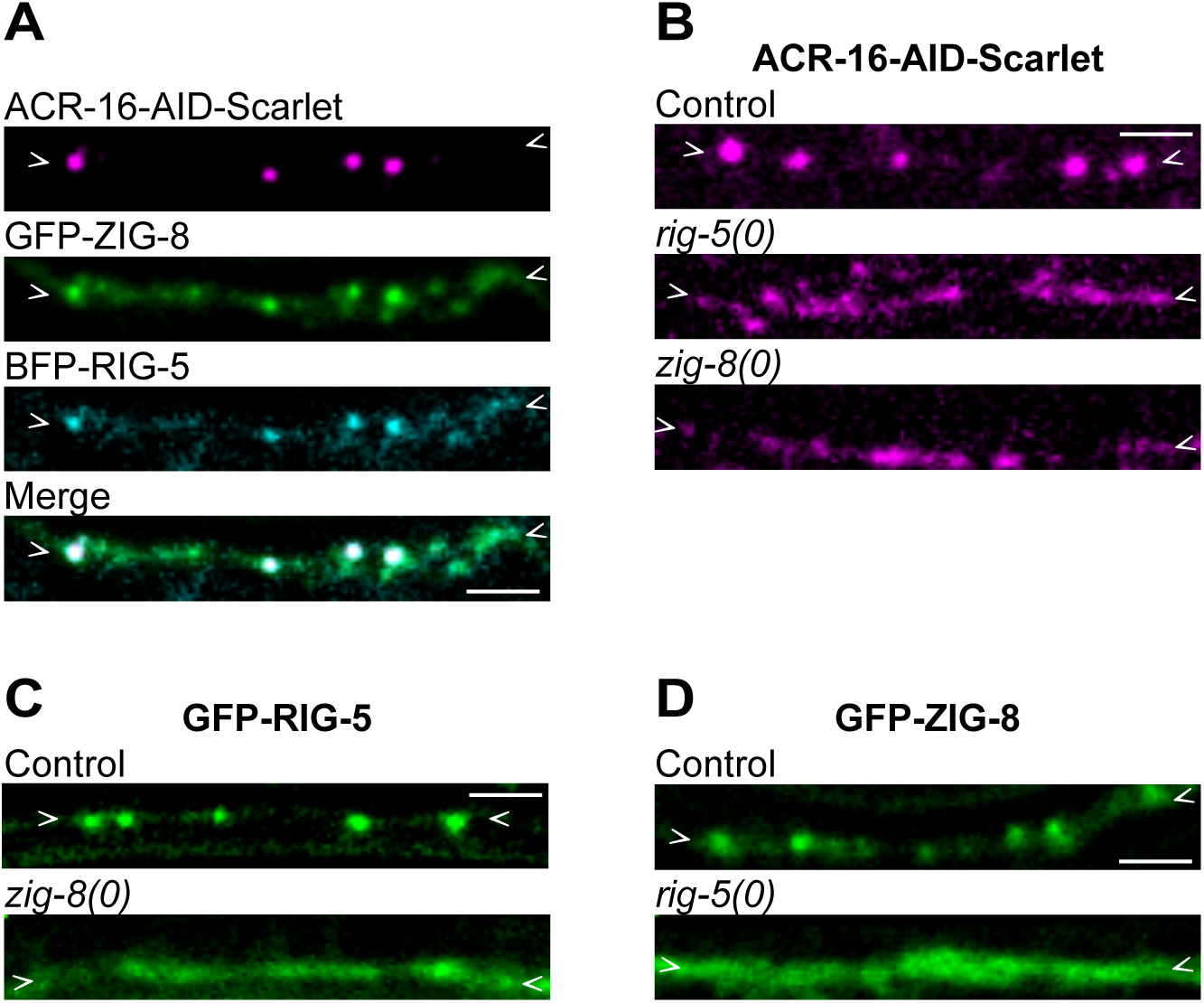
RIG-5 and ZIG-8 control ACR-16 receptor clustering in L1 larvae. **(A)** Knock-in reporters of BFP-RIG-5 and GFP-ZIG-8 show strong colocalization with neuronal ACR-16-AID-Scarlet. **(B)** ACR-16-AID-Scarlet fluorescence shows clustering defects in neurons of the *rig-5(0)* and *zig-8(0)* mutants. **(C)** GFP-RIG-5 observed in a control strain and in the *zig-8(0)* mutant. **(D)** GFP-ZIG-8 observed in a control strain and in the *rig-5(0)* mutant. In this figure, all spinning disk microscopy images were taken in newly born L1 animals. For B, C and D, fluorescence intensities were enhanced in the *rig-5(0)* and *zig-8(0)* mutants. Scale bars: 2 µm.

**Supplementary Figure 5.**
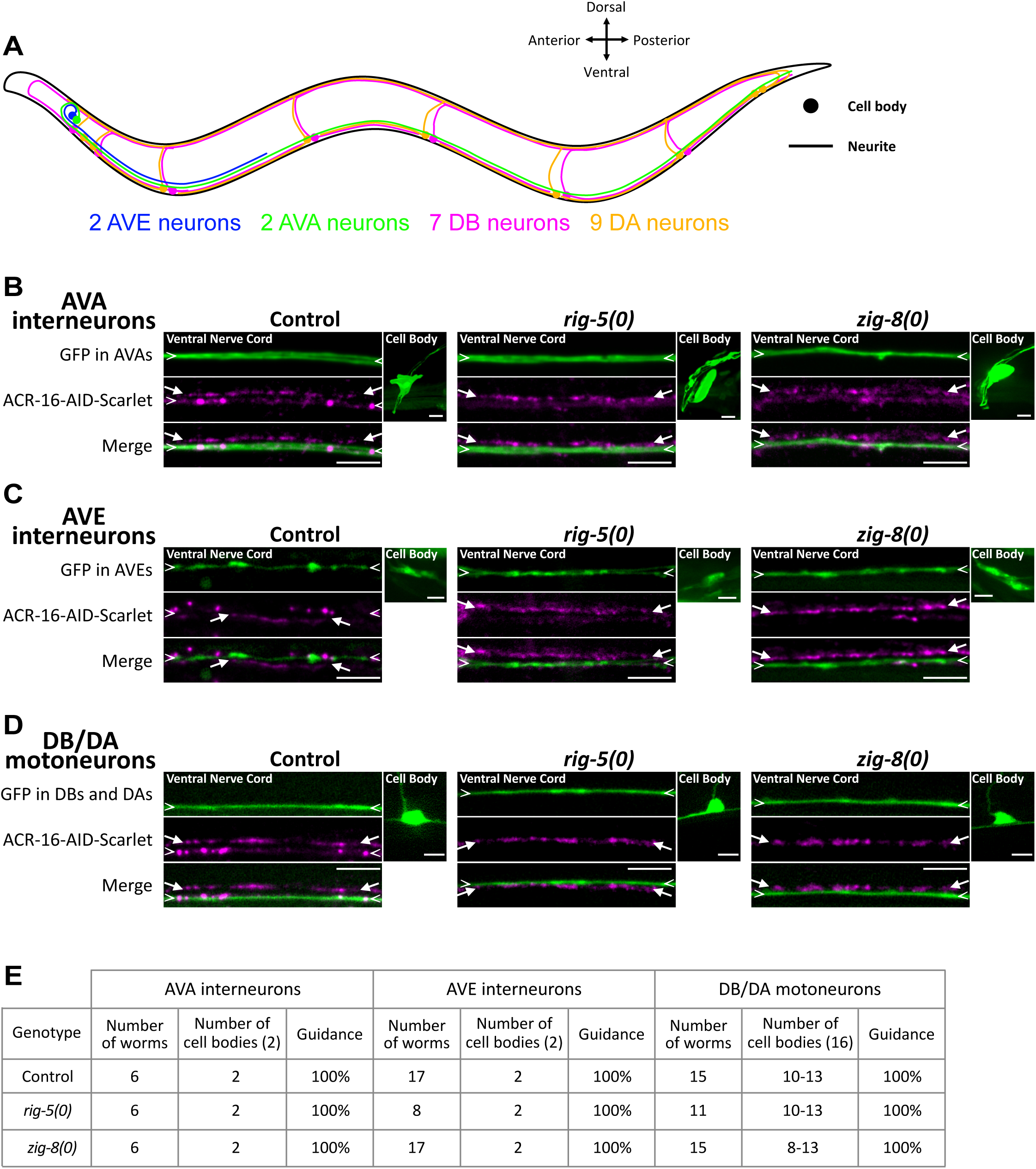
*rig-5* and *zig-8* mutations do not alter neuronal specification, migration or axon guidance of AVA, AVE and DB neurons. **(A)** Schematic of cell bodies and neurites of AVE and AVA neurons, and DB-DA-types motoneurons (comprising the 7 DB-type and 9 DA-type neurons) in an adult *C. elegans*. **(B-D)** Assessment of the presence and position of cell bodies and neurites using cytoplasmic GFP specifically expressed in AVA (B), AVE (C) neurons and DB-DA-types motoneurons (D) in control backgrounds and in *rig-5(0)* and *zig-8(0)* mutants. Scale bars: 5 µm. **(E)** Summary table.

**Supplementary Figure 6.**
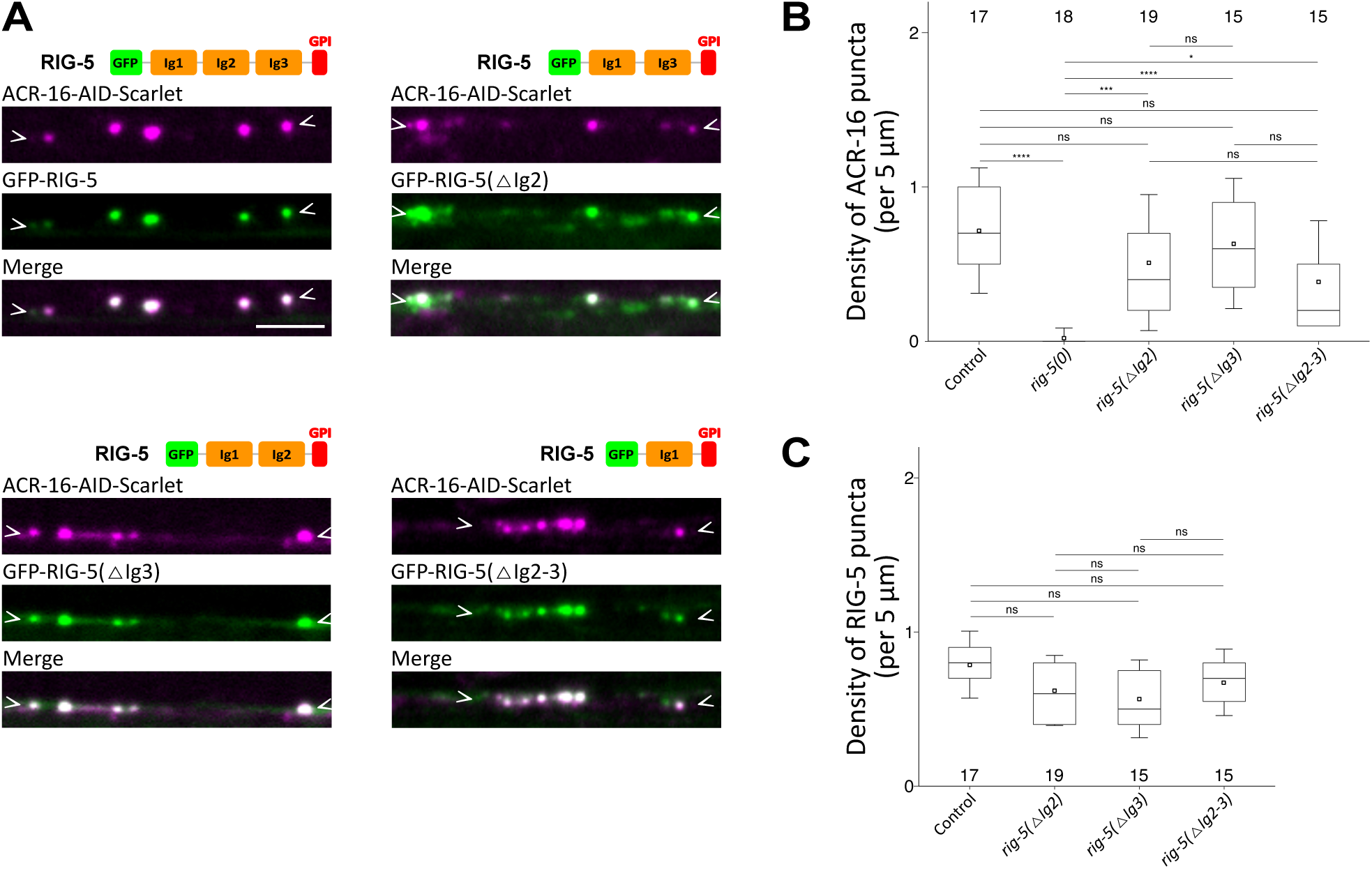
RIG-5 Ig2 and Ig3 domains are dispensable for ACR-16 receptor clustering. **(A)** ACR-16-AID-Scarlet was observed in control, *gfp::rig-5*, *gfp::rig-5(Δ1Ig2)*, *gfp::rig-5(Δ1Ig3)* and *gfp::rig-5(Δ1Ig2-3)* following auxin-induced degradation in muscle cells. A schematic of RIG-5 functional domains is shown for each knock-in condition. Scale bar: 5 µm. **(B-C)** Quantification of ACR-16 (B) and RIG-5 (C) puncta density. Data are presented as boxplots showing lower and upper quartiles (box), mean (square), median (center line) and standard deviation (whiskers); the number of worms is indicated for each condition; Kruskal-Wallis and Dunn’s test (B-C); ns: non-significant, *<0.05, ***<0.0005, ****<0.00005.

**Supplementary Figure 7.**
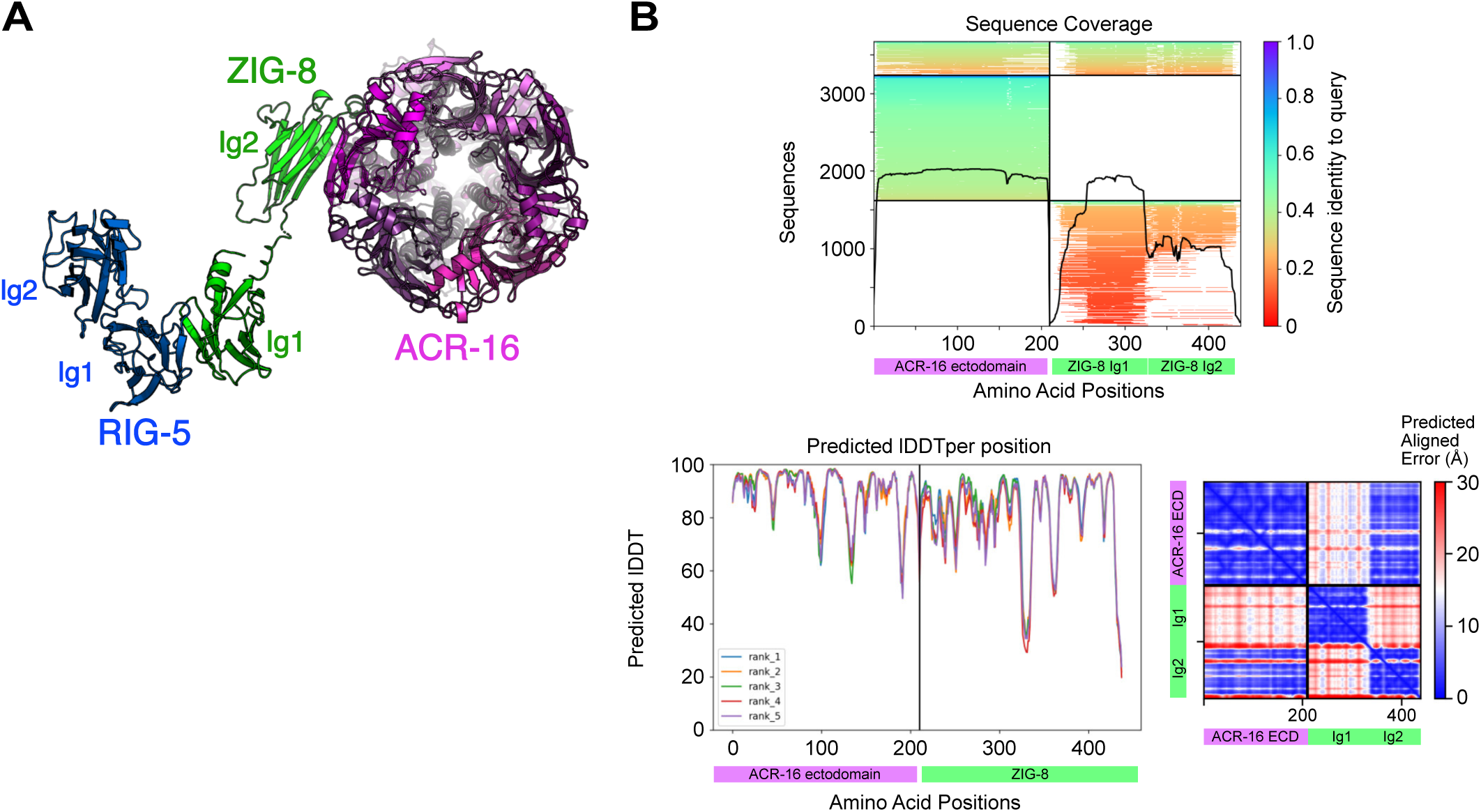
Structure prediction of the RIG-5—ZIG-8—ACR-16 complex. **(A)** Structural model for the composite RIG-5—ZIG-8—ACR-16 complex with a top view showing the channel pore. **(B)** Colabfold plots for multiple sequence alignment sequence coverage, predicted local distance difference test (pLDDT), and predicted aligned error for the ACR-16 ectodomain complex with ZIG-8.

**Supplementary Figure 8.**
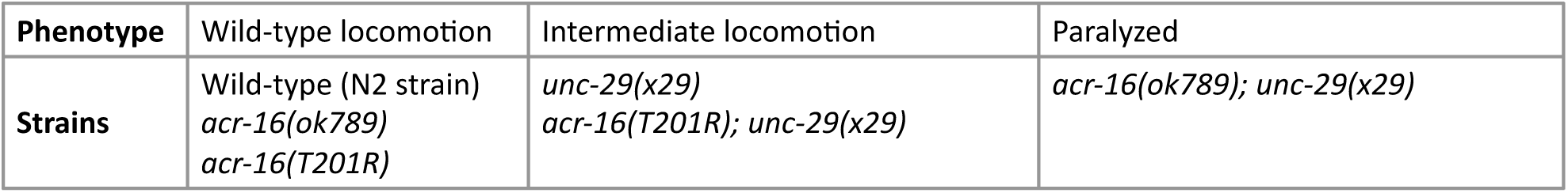
The T201R mutation does not impair the function of ACR-16 receptors. Locomotion test of control, *acr-16(ok789), unc-29(x29), acr-16(T201R)* single mutants, as well as *acr-16(ok789); unc-29(x29)* and *acr-16(T201R); unc-29(x29)* double mutants.

**Supplementary Table 1. Strain list.**

**Supplementary Table 2. Allele list.**

**Supplementary Table 3. List of guides used for CRISPR/Cas9 gene editing.**

**Supplementary Table 4. Plasmid list.**

**Supplementary Table 5. List of primers used to amplify promoters.**

**Supplementary Table 6. Composition of the injection mix used to generate extrachromosomal array lines.**

## Notes

### Competing Interest Statement

The authors have declared no competing interest.

